# Comparative Analysis of Molecular Pathogenic Mechanisms and Antiviral Development Targeting Old and New World Hantaviruses

**DOI:** 10.1101/2023.08.04.552083

**Authors:** Arjit Vijey Jeyachandran, Joseph Ignatius Irudayam, Swati Dubey, Nikhil Chakravarty, Bindu Konda, Aayushi Shah, Baolong Su, Cheng Wang, Qi Cui, Kevin J. Williams, Sonal Srikanth, Yanhong Shi, Arjun Deb, Robert Damoiseaux, Barry R. Stripp, Arunachalam Ramaiah, Vaithilingaraja Arumugaswami

## Abstract

**Background:** Hantaviruses – dichotomized into New World (i.e. Andes virus, ANDV; Sin Nombre virus, SNV) and Old-World viruses (i.e. Hantaan virus, HTNV) – are zoonotic viruses transmitted from rodents to humans. Currently, no FDA-approved vaccines against hantaviruses exist. Given the recent breakthrough to human-human transmission by the ANDV, an essential step is to establish an effective pandemic preparedness infrastructure to rapidly identify cell tropism, infective potential, and effective therapeutic agents through systematic investigation.

**Methods:** We established human cell model systems in lung (airway and distal lung epithelial cells), heart (pluripotent stem cell-derived (PSC-) cardiomyocytes), and brain (PSC-astrocytes) cell types and subsequently evaluated ANDV, HTNV and SNV tropisms. Transcriptomic, lipidomic and bioinformatic data analyses were performed to identify the molecular pathogenic mechanisms of viruses in different cell types. This cell-based infection system was utilized to establish a drug testing platform and pharmacogenomic comparisons.

**Results:** ANDV showed broad tropism for all cell types assessed. HTNV replication was predominantly observed in heart and brain cells. ANDV efficiently replicated in human and mouse 3D distal lung organoids. Transcriptomic analysis showed that ANDV infection resulted in pronounced inflammatory response and downregulation of cholesterol biosynthesis pathway in lung cells. Lipidomic profiling revealed that ANDV-infected cells showed reduced level of cholesterol esters and triglycerides. Further analysis of pathway-based molecular signatures showed that, compared to SNV and HTNV, ANDV infection caused drastic lung cell injury responses. A selective drug screening identified STING agonists, nucleoside analogues and plant-derived compounds that inhibited ANDV viral infection and rescued cellular metabolism. In line with experimental results, transcriptome data shows that the least number of total and unique differentially expressed genes were identified in urolithin B- and favipiravir-treated cells, confirming the higher efficiency of these two drugs in inhibiting ANDV, resulting in host cell ability to balance gene expression to establish proper cell functioning.

**Conclusions:** Overall, our study describes advanced human PSC-derived model systems and systems-level transcriptomics and lipidomic data to better understand Old and New World hantaviral tropism, as well as drug candidates that can be further assessed for potential rapid deployment in the event of a pandemic.

## INTRODUCTION

Hantaviruses have been major human pathogen of concern for several decades and are classified as Old World and New World hantaviruses. Hantaviruses first came to the world’s attention during the Korean War in the 1950s. Over 3000 troops fell ill with Korean hemorrhagic fever, now referred to as hemorrhagic fever with renal syndrome (HFRS)^1^ – the etiological agent of which is now known to be Old World Hantaan virus (HTNV), named after the Hantan River located near the outbreak sites. Hantaviruses are mostly transmitted by rodent and insectivore vectors, often mice and rats, via aerosolized excrement – a distinguishing factor from the other members of the family *Hantaviridae*, which are transmitted via arthropod vectors^2^. HTNV is transmitted by a rodent species *Apodemus agrarius*, which is primarily found within Central Europe, Russia, Korea and China^3, 4^. Old World Sin Nombre virus (SNV) is carried by *Peromyscus maniculatus*, which inhabit the United States and Canada^5, 6^. New World Andes virus (ANDV) is carried by *Oligozomys longicaudatus*, which reside in Argentina along the Andes mountains^3^. Most hantavirus species have not been detected to be transmitted between humans, thus their main mode of transmission is through animal reservoir. Several pathogenic viral species have been identified in recent decades, causing HFRS [e.g., Puumala virus (PUUV)^7^, HTNV] or hantavirus cardiopulmonary syndrome (HCPS) [e.g., SNV^5^, ANDV^8^]. Depending on the infecting strain, the mortality rate can differ across a range of 1% to 40%^9^. ANDV has shown evidence of human-human transmission, making it a viral disease of major concern^10, 11^. The scope of this study mainly focuses on the Old World HTNV and the New World SNV and ANDV, as these viral species present significant danger to the world population.

Hantaviruses contain an 11-19kb tri-segmented, negative-sense, single-stranded RNA genome split into a small (S), medium (M), and large (L) segment within an envelope covered in viral glycoproteins^12^. Hantavirus Gn/Gc glycoproteins are the only viral proteins exposed on the surface of virions, which facilitate virus attachment and entry to the cells. Although the overall architecture of Gn/Gc complex is conserved among hantaviruses, Gn is less conserved than Gc and is proposed to bind to cell-surface receptors during viral entry. Due to their variability in Gn sequence, distinct hantavirus clades are likely to use different attachment factors and/or receptors^13–15^. Hantaviruses are particularly dangerous due to their ability to infect a wide variety of cells across several different vital systems – namely, the renal, pulmonary, nervous, and cardiac systems^9^. Evidence has been presented that hantaviruses readily infect endothelial^16^ and mononuclear^17–24^ cells through a less-understood mechanism and use these cells as vehicles to spread throughout the body to nearly all major organs^16^.

Despite research efforts, there have been no effective prophylactic or therapeutic antiviral agents approved targeting hantaviruses. Virus-like particle, inactivated, recombinant, viral vector, and nucleic acid-based vaccine candidates have been developed and are either currently being tested or undergoing clinical trials with varying ability to prevent infection by HTNV, PUUV, SEOV, ANDV, and DOBV^25^. The drug vandetanib showed semi-prophylactic ability in ANDV-infected Syrian hamsters, delaying lethality and increasing total survival by 23% when administered five days prior to ANDV challenge^26^. Several antiviral drug candidates have also been investigated for their effects in treating hantavirus disease. Ribavirin, though promising in mouse and Syrian hamster models^27–30^, was generally not seen to be effective in humans during clinical trials^31–33^. Other drug candidates have been tested in animal models, showing some efficacy in treating hantavirus infection, such as lactoferrin^34^, ETAR (1-beta-d-ribofuranosyl-3-ethynyl-[1,2,4] triazole)^35^, and favipiravir^36^.

Moreover, the mechanism of hantaviral disease pathophysiology is still poorly understood. Thus, we aim to assess differences in viral tropism between Old and New World hantaviruses. In this study, we developed advanced human stem cell-derived cellular and organoid systems to model and investigate differences in pathogenic processes at the molecular level, as well as cell injury mechanisms.

## RESULTS

### New World ANDV Displays Broad Cell Tropism

In order to understand the cell tropism and cell injury mechanism of phylogenetically related Old and New World hantaviruses (Figure 1A, Table S1), we set out to establish human cell culture model systems. We used Old World HTNV (Fojnica strain) as well as New World ANDV (Chile-9717869 strain) and SNV (SNV-77734) for infection studies, as illustrated in Figure 1B. All three viruses were amplified in Vero E6 cells and replicated at similar levels (Figure 1C). Virus replication was quantified using strain-specific primer sets using sensitive RT-qPCR (Figure 1C; refer to Methods).

**Figure 1.**
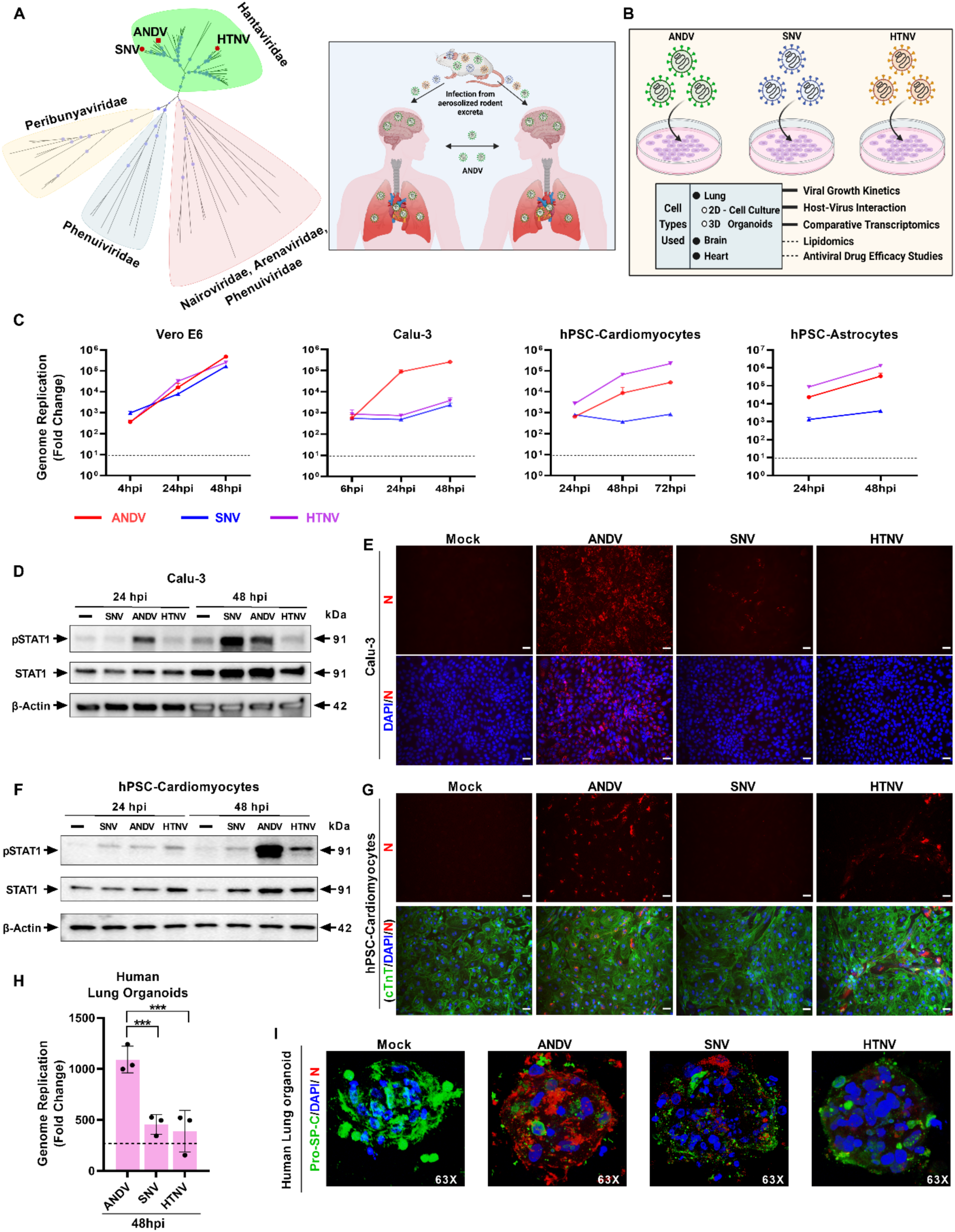
Cell tropism of various Hantaviruses. **A)** Phylogenetic analysis of aligned and sequenced M segment of viral sequences (n = 68) from *Hantaviridae, Phenuiviridae, Nairoviridae, Arenaviridae,* and *Peribunyaviridae* families of the *Bunyavirales* order. In the *Hantaviridae* cluster: SNV = circle; ANDV = square; and HTNV = star. The adjacent diagram presents the transmission modes of these three viruses. **B)** Schematic outline of cell types used, and methodologies employed in this study. **C)** The graphs demonstrate the differing levels of viral genome replication of hantaviruses in indicated cell types. **D, F)** Western blot analysis of cells infected with hantaviruses. **E, G)** Immunofluorescent analysis detects viral nucleocapsid (red) and cardiac troponin (green) in hantavirus-infected cells at 48 hours post-infection (hpi). Scale bar: 25μm. **H)** The graph shows the levels of viral genome replication in human distal lung 3D organoids at 48 hpi. **I)** Confocal microscopy images of human distal lung 3D organoids depict lung marker pro-SP-C (green) and hantaviral N protein (red). Scale bar: 50μm. Quantitative data are presented as mean ± standard deviation. Statistical analysis was performed using ANOVA, followed by Tukey’s post hoc test (*, P < 0.05; **, P < 0.01; ***, P < 0.001). Representative data presented from two or more experimental repeats.

Since the New World hantaviruses cause cardiopulmonary diseases, we first examined the susceptibility of lung and heart cell types. Calu-3 lung cells were infected with ANDV, SNV and HTNV viruses at an MOI of 0.1. At 6, 24, and 48 hpi, cellular RNA was collected, and viral load were assessed by RT-qPCR. Though we observed significant replication of all three viral species in lung cells, ANDV had over a 100-fold higher replication rate than SNV or HTNV (Figure 1C). Western blot analysis revealed that ANDV-infected lung cells had type I IFN signaling pathway activation, as determined by phosphorylation of STAT1 (Figure 1D). Another New World cardiopulmonary disease-causing virus, SNV, also activated this pathway at 48 hpi. Interestingly, STAT1 phosphorylation was below detectable levels in HTNV-infected cells (Figure 1D). Immunofluorescence analysis (IFA) of these cells using antibody detecting viral nucleocapsid (N) confirmed that ANDV infected most of the cells (Figure 1E).

It has been well established that HTNV and ANDV can readily infect human endothelial cells^25–28^. As such, we focused our effort on better understanding hantaviral infection in heart cells, using human pluripotent stem cell-derived cardiomyocytes (hPSC-CMs). In this cell type, both ANDV and HTNV replicated at a higher level, whereas SNV established a low-grade infection (Figure 1C). This finding was further confirmed by IFA and Western blot analysis (Figures 1F, 1G). In SNV-infected cells, the N protein was not detected (Figure 1G). Subsequently, to assess the host innate immune and inflammatory responses, we evaluated the gene expression levels of OAS1, IFN-λ, IFN-β, and IL1-β in both Calu-3 cells and hPSC-CMs (Figure S1). Our observations revealed a significant upregulation of these genes upon ANDV infection over time in both cell types (Figure S1A). Interestingly, in hPSC-CMs, HTNV had a higher viral load than ANDV. At the functional level, ANDV-infected hPSC-CM organoids had significantly reduced cell-beat contractions (Figure S1 and Supplementary Videos). Moreover, ANDV-infected cells demonstrated increased levels of immune gene activation, suggesting different levels in virulence between these New and Old World viruses.

Hantaviruses have also been shown to cause neurological symptoms and disease^29–32^. As such, we further investigated the infectivity of these viruses in human astrocytes, a major brain cell type (Figure S1B). Mouse model studies using HTNV showed brain infection and resulting neurological signs, such as paralysis^33, 34^. We observed that both ANDV and HTNV established active infection in hPSC-derived astrocytes (hPSC-astrocytes) (Figure 1C, Figure S1B), suggesting that ANDV can also cause neurological disorders in human.

After assessing differential tropism in 2D cell culture, we evaluated viral replication using a primary human distal 3D lung organoid model (Figures 1H and 1I). Through this model system^35^, we observed that ANDV replicated at a higher level compared to HTNV and SNV, which was similar to observed responses in Calu-3 cells (Figure 1C). Presence of alveolar type 2 (AT2) cells within distal lung organoids was confirmed by staining for the AT2 cell marker, Pro-SP-C (Figure 1I). Because rodents are the natural reservoir for hantaviruses, we subsequently evaluated the replication of these viruses in a laboratory mouse distal lung organoid system. In this system, we surprisingly observed that both ANDV and HTNV replicated at similar levels (Figures S1C and S1D). In general, the SNV strain that we used exhibited poor replication efficiency in all tested cell systems. Our observations regarding SNV inefficiency in replication can partially be explained by the lack of disease phenotype in animal models^36–39^. Based on these results, we conclude that the New World ANDV exhibits broader tissue tropism.

### Comparative Transcriptomics Analysis of Old and New World Hantaviruses

We investigated host cellular responses and associated dysregulated pathways to Old and New World hantaviral infection of lung, heart, and brain cell types using systems-level transcriptomics (Figure 2A). We used the Calu-3 cells, hPSC-CMs and hPSC-astrocytes infected with each virus (MOI 0.1). At 48 hpi, the cells were harvested for RNA sequencing. A total RNA sequencing analysis of virus-infected and uninfected cell types showed that ANDV-infected cell types had a larger number of differentially expressed genes (DEGs) compared to HTNV and SNV (Figure 2A). HTNV- and SNV-infected cells show similar transcriptional responses, though significantly different from ANDV-infected cells. For instance, the expression of host genes in ANDV-infected lung cells was increased about 73- and 27-fold compared to HTNV- and SNV-infected cells, respectively, indicating viral lineage-specific pathogenesis mechanisms of hantaviruses and associated host cell responses (Figures 2A-2C). The ANDV-infected hPSC-astrocytes also exhibited increased transcriptional responses (7-fold to HTNV; 11-fold to SNV). However, no significant transcriptional differences were observed between these viruses following infection of hPSC-CMs, reflecting cell-specific differences in tropism. These data also illustrate that, while lung followed by brain cells are more susceptible to ANDV, heart cells are likely susceptible to all three viral lineages. HTNV- and SNV-infected cells exhibited similar gene expression patterns, indicating that infection with either of these two viruses had a minimal effect irrespective of cell type at 48 hpi. Comparison of ANDV- with HTNV- or SNV-infected cells displayed a higher number of upregulated genes [Calu-3: 1003 (ANDV vs. HTNV) and 915 (ANDV vs. SNV); astrocytes 370 (ANDV vs. HTNV) and 558 (ANDV vs. SNV); hPSC-CMs 188 (ANDV vs. HTNV) and 205 (ANDV vs. SNV)], indicating that the pathogenic mechanism of virulent ANDV is different relative to the other two viruses (Figures 2B and 2C). Comparison of HTNV- with SNV-infected cells revealed no noticeable difference in transcriptional responses in all host cell types, suggesting that the properties that contribute to the virulence of these two viruses are similar despite representing Old World and New World lineages, respectively. It is important to consider that these two strains are similar in their zoonotic transmission, lacking the ability to transmit between humans (Figure 1A). Overall, the host transcriptional responses to each of three hantaviruses were differential in nature, showing a range of only 19-79 commonly downregulated and 18-78 upregulated genes, suggesting that genes involved in a few biological functions are commonly dysregulated by these viruses. Reflecting the *in vitro* experimental data, transcriptome data from three cell types also revealed that ANDV infection had the largest number of uniquely downregulated (Calu-3: 431; astrocytes: 200; hPSC-CM: 17) and upregulated (Calu-3: 969; astrocytes: 649; hPSC-CM: 222) genes (Figure 2C). These genes may be viral lineage-specific responses induced by different cell types, implying the extent of organ susceptibility to the same virus.

**Figure 2.**
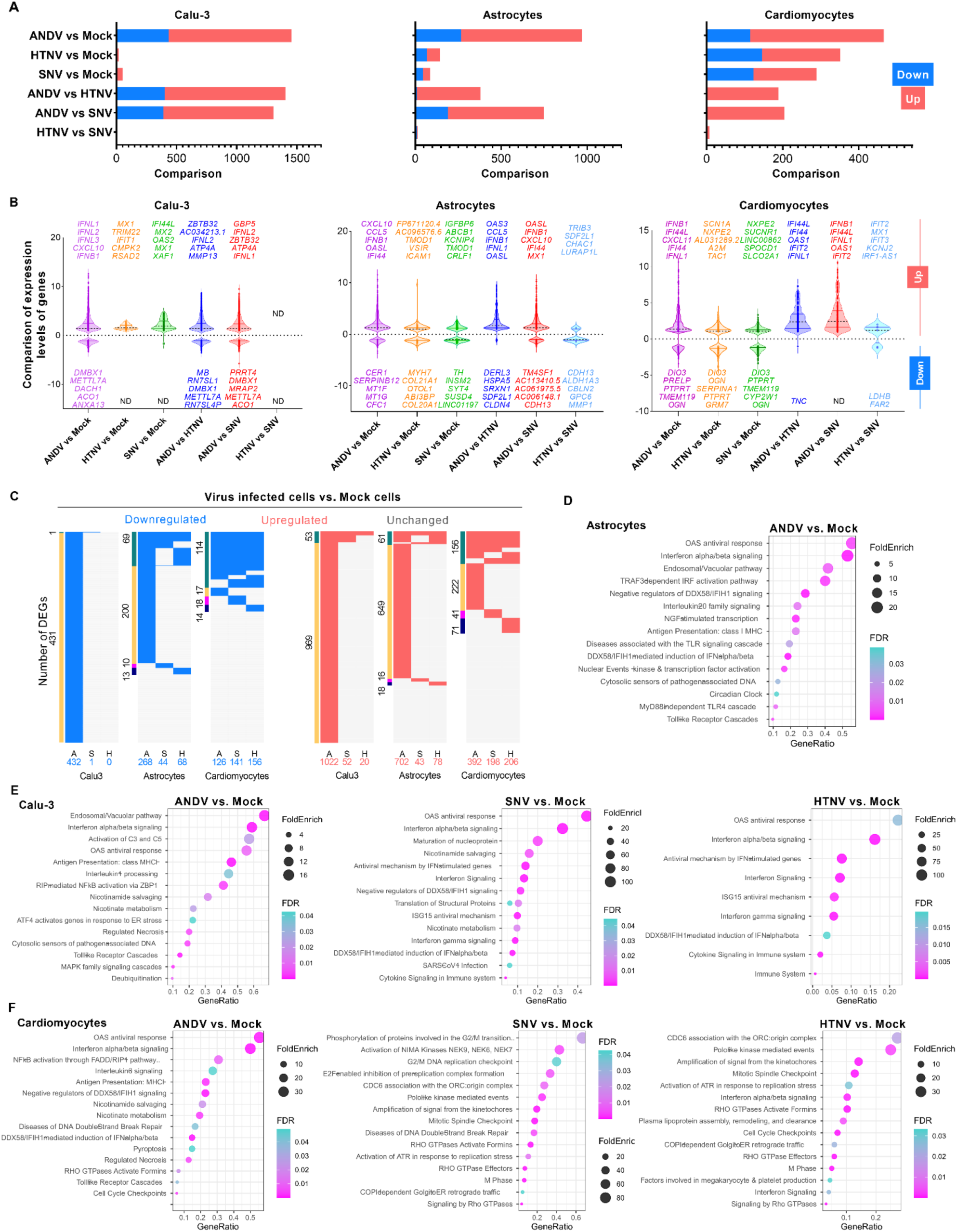
Comparative transcriptomic analysis of hantaviruses. **A)** Bar graph shows the total number of down- and up-regulated genes in Calu-3, hPSC-astrocyte and hPSC-CM cells infected with indicated viruses. **B)** Violin plots show comparison of expression levels (log2 Fold Change) of differentially expressed genes (DEGs) in various samples. **C)** Comparison of down- and up- regulated DEGs in cells infected with indicated viruses (A, ANDV; S, SNV; H, HTNV). The side bars show DEGs group (green: DEGs shared by at least two datasets; others: exclusive to each dataset). **D)** Dot plot shows the up-regulated pathways in ANDV-infected hPSC-astrocytes. **E, F)** Dot plots represent the up-regulated pathways in infected Calu-3 and hPSC-CMs, respectively.

We identified 316 commonly upregulated genes (*FDR* 0.01) across three human cell types infected with ANDV (Figure 2C, Table S2). Of these 316 genes, 20 were highly upregulated (>5 Log2FoldChange) in these cell types, most of which were related to innate immune and inflammatory chemokines (Table S2). Similarly, these human cell types infected with HTNV had six commonly upregulated genes (*TAP1, DTX3L, DUSP1, PARP9, IFI16, REC8*), whereas SNV had three (*PLSCR1, DTX3L, IFI16*). Genes *DTX3L* and *IFI16* were found to be commonly induced across all three hantaviral infections. *PLSCR1* transcriptional induction was shared by both New World hantaviruses. These upregulated biomarkers may reflect the differential susceptibility and host responses of the tested cell types to these viruses.

Among the twelve viral entry receptor genes, six were transcriptionally stimulated by hantaviral infection in the three cell types studied (Table S3). We identified ICAM1 to be commonly upregulated in all three cell types upon ANDV infection whereas it was only upregulated in two cell types (hPSC-CMs and -astrocytes) upon HTNV infection and only in hPSC-astrocytes upon SNV infection. It should also be noted that ICAM1 was upregulated during inflammatory response. GRIK1 was downregulated in hPSC-CMs and -astrocytes infected with hantaviruses. ANDV infection showed upregulation of cell type-specific entry genes (Calu-3: *CD55* and *ITGB3*; hPSC-astrocytes: *MERTK* and *VEGFA*). HTNV infection upregulated the *HAVCR2* gene in hPSC-astrocytes. Further exploiting these viral entry receptors and their interactions with viruses would be an attractive target for new antiviral therapeutics and vaccine technologies.

Pathway and Gene Ontology enrichment analyses of infected Calu-3 cells showed upregulation of immune pathways, including OAS antiviral response, IFN-α/β signaling, IFN-γ signaling, and ISG15 antiviral mechanism by all three viruses. However, cell death mechanisms, including programmed cell death, death receptor signaling and necrosis (Figure 2E), and immune pathways such as many interleukin signaling and TLR cascades, and the MyD88-independent cascade were upregulated only by ANDV. These molecular signatures suggest that, compared to SNV and HTNV viruses, ANDV infection drastically damages lung cells. In hPSC-astrocytes, no pathways were upregulated by SNV and HTNV infection while many immune pathways were upregulated by ANDV (Figure 2D). Pathway analysis of hPSC-CMs showed that upregulation of cell cycle pathway genes was common across all three viruses while cell repair and related pathways were only upregulated in SNV and HTNV (Figure 2F). In contrast, ANDV infection upregulated many immune and inflammatory, as well as cell death and necrosis, pathways in hPSC-CMs (Figures S2A – S2D). Taken together, our comprehensive transcriptome analysis revealed that i) enriched pathophysiological molecular changes occurred in all three cell types infected with ANDV compared to the other two viruses, and ii) ANDV infection leads to lung and heart cell injury and death. These findings imply that we may expect outbreaks in the future by rapidly spreading sub-strains of ANDV to cause serious global health crisis.

### Cholesterol Pathway is Downregulated Upon ANDV Infection

Upon identifying these transcriptional pathways, we set out to identify metabolic pathways dysregulated during ANDV infection. Our transcriptomic analysis indicated that ANDV infection significantly downregulated the cholesterol synthesis pathways in Calu-3 cells and hPSC-astrocytes (Figure 3A, Figure S3A). Genes controlling the key steps in cholesterol biosynthesis, such as *FDPS, ACAT2, MVD, LSS* and *FDFT1*, were transcriptionally commonly suppressed upon viral infection at 48 hpi (Figure 3B and 3C, Figure S3B). This finding is surprising because ANDV has been reported to require cholesterol for viral entry and replication^40^. In the context of host-virus interactions, this observation perhaps could be due to host antiviral response to deplete cholesterol in infected cells to restrict virus production. However, this depletion has the potential to damage cellular processes and thus play a role in hantaviral pathogenesis.

**Figure 3.**
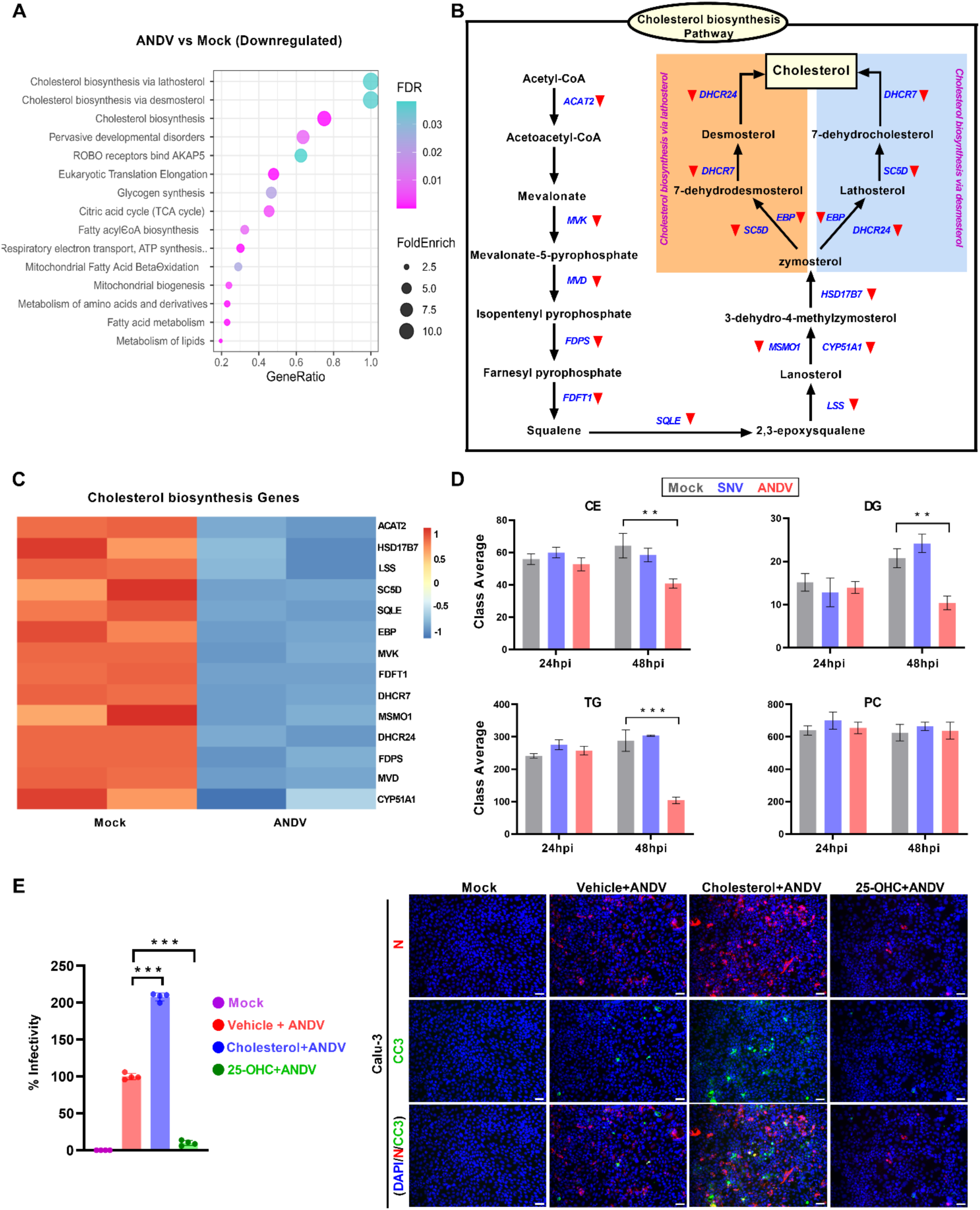
Cholesterol pathway dysregulation in ANDV-infected Calu-3 cells. **A)** The dot plot illustrates downregulated pathways, including cholesterol synthesis in ANDV infected cells relative to uninfected cells at 48 hpi. **B)** The diagram illustrates the downregulated genes in cholesterol synthetic pathway. **C)** Heatmap shows Z scores as expression levels of the genes involved in cholesterol biosynthesis pathway. Blue and red colors represent downregulated and upregulated genes, respectively. **D)** Graphs indicate levels of lipid molecule subclasses at 24 hpi and 48 hpi. **E)** Graph shows the percent infectivity of ANDV at 48 hpi in indicated treatment conditions. Immunofluorescent analysis of ANDV-infected cells supplemented with cholesterol or 25-OHC at 48 hpi. Nucleocapsid = red; Cleaved caspase-3 = green. Scale bar: 25μm. Quantitative data are presented as mean ± standard deviation. Statistical analysis was performed using ANOVA, followed by Tukey’s post hoc test (*, P < 0.05; **, P < 0.01; ***, P < 0.001).

To verify these transcriptional changes in cholesterol and lipid metabolites within infected cells, we carried out a lipidomics study. We observed that ANDV-infected Calu-3 cells had significantly lower levels of cholesterol esters, triglycerides, and diglycerides at 48 hpi (Figure 3D). SNV infection of Calu-3 cells did not affect the levels of these metabolites. Phosphotidylcholine (PC) and phosphotidylethanolamine (PE) levels were not affected in both ANDV- and SNV-infected cells. However, levels of their intermediate metabolites lysophosphotidylcholine (LPC) and lysophosphotidylethanolamine (LPE) were increased upon ANDV infection (Figure S3C). We also observed reductions in triglyceride (TG) bonds and TG carbons in ANDV-infected cells (Figure S3D). Therefore, this reduction in TG bonds and TG carbons likely confirms viral impact on the host cell lipid metabolism by dysregulating the process involved in triglyceride synthesis.

Since cholesterol is important for ANDV replication, we performed a study where cholesterol was supplemented in cell media of ANDV-infected Calu-3 cells. We observed that supplementary cholesterol significantly enhanced ANDV infection (Figure 3E). This experiment confirmed that the depletion of cholesterol pathway-associated genes and metabolites observed at the transcriptional and metabolomic levels is likely induced by host antiviral response to ANDV infection. Furthermore, we observed that the gene *CH25H*, which encodes for the IFN-stimulated enzyme cholesterol 25-hydroxylase, was upregulated in ANDV-infected cells. This gene produced the antiviral metabolite 25-hydroxycholesterol (25-OHC). As such, we assessed the potential anti-hantaviral activity of 25-OHC. The addition of 25-OHC inhibited the replication of the virus in the infected cells (Figure 3E). These results suggest that ANDV infection leads to dysregulation of the cholesterol synthesis pathway, potentially causing disturbances in the metabolic processes of the host cell.

### Establishing a Drug Screening Platform to Identify Potential Antivirals Against ANDV

Because of its known ability for human-human transmission, as well as our observation of higher virulence in a broader set of cell types, we set out to identify potential antiviral agents targeting ANDV. For this purpose, we utilized the known antiviral biologic IFN-β, as well as the STING pathway agonists diABZI and cAIMP, to establish a drug testing platform^53^ (Figure S4A). Calu-3 cells were pre-treated with drug compounds at 24h prior to infection and cells were fixed at 48 hpi. RNA samples were then collected to assess viral load. As a negative control, we used the protease inhibitor compound GC376, which has been shown to be effective against viruses encoding proteases, such as SARS-CoV-2^54^. We observed that both IFN-β and STING agonists effectively inhibited ANDV replication (Figures S4A–S4C). As expected, GC376 did not show inhibition against ANDV, which does not encode for a protease.

Once this drug testing platform was established, we subsequently tested the antiviral effectiveness of nucleoside analogs (favipiravir^55^ and 6-azauridine [6AZA^56^]), as well as herbal compounds (silymarin^57^ and urolithin B^58^), shown to have antioxidant properties (Table S4). We observed that these compounds significantly inhibited ANDV genome replication and infection at non-cytotoxic dose levels (Figure 4A and 4B, Figure S4D). To understand the mode of action of these antiviral compounds, we utilized RNA-Seq methodologies to profile the transcriptomic changes. We found that host transcriptional response was differential in drug-treated cells, suggesting varying levels of ANDV inhibition in combination with the drug-specific host effect (Figures 4C-4E). This differential response was confirmed by the extreme coordinates on the principal-component analysis (PCA) plot for the drug conditions tested (Figure 4C). Cells treated with 6AZA and silymarin showed similar transcriptional response profiles, which varied from those seen in urolithin B- and favipiravir-treated cells when compared to vehicle-treated cells (Figures 4D and 4E). There was significant overlap in DEGs across drug-treated infected cells, with the majority of the 799 commonly upregulated and 697 commonly downregulated genes being shared across two or more drug conditions when compared to uninfected (mock) cells (Figure 4F). Among these DEGs, only 27 downregulated and 40 upregulated genes were broadly expressed across the four drug-treated conditions, suggesting that genes involved in only certain molecular functions are commonly induced by four drugs. However, drug-treated infected cells had a range of 6 to 4,323 exclusive DEGs that could possibly be drug-specific elements contributing to the antiviral process and cell recovery when compared to mock. In line with experimental results, it is noteworthy to observe that the least number of total and unique DEGs were identified in the urolithin B- and favipiravir-treated cells, confirming the higher efficiency of these two drugs in inhibiting ANDV, resulting in host cell ability to balance gene expression to establish proper cell functioning (Figures 4D-F, Figure S4D). Host unique gene expression in nucleoside analog 6AZA-treated infected cells was increased 721- and 330-fold as compared to favipiravir- and urolithin B-treated infected cells, respectively. This finding implies that, in addition to arresting the viral replication, 6AZA may also be involved in dysregulating thousands of host genes and many pathways, as well as the transcription process^59^, which can manifest as adverse or detrimental effects. Pathway and Gene Ontology enrichment analyses of drug-treated infected cells showed upregulation of pathways involved in cell repair, cellular responses, and mitochondrial biogenesis and metabolisms, including glycogen metabolism, amino acids and derivates, metabolisms of glucose, fatty acid, carbohydrates, proteins, vitamins and cofactors and lipids, as compared to vehicle-treated infected cells (Figure 4G, Table S4). In contrast, 6AZA further downregulated most of these pathways compared to vehicle-treated infected cells. We did observe that many immune, signaling and cell death pathways, including apoptosis, death receptor signaling, necrosis, IFN signaling, NF-κB, interleukin-10 signaling, and cytokine signaling, were commonly inhibited or downregulated in cells treated with any of the four drugs. This is reflective of reduced viral replication because of drug treatment.

**Figure 4.**
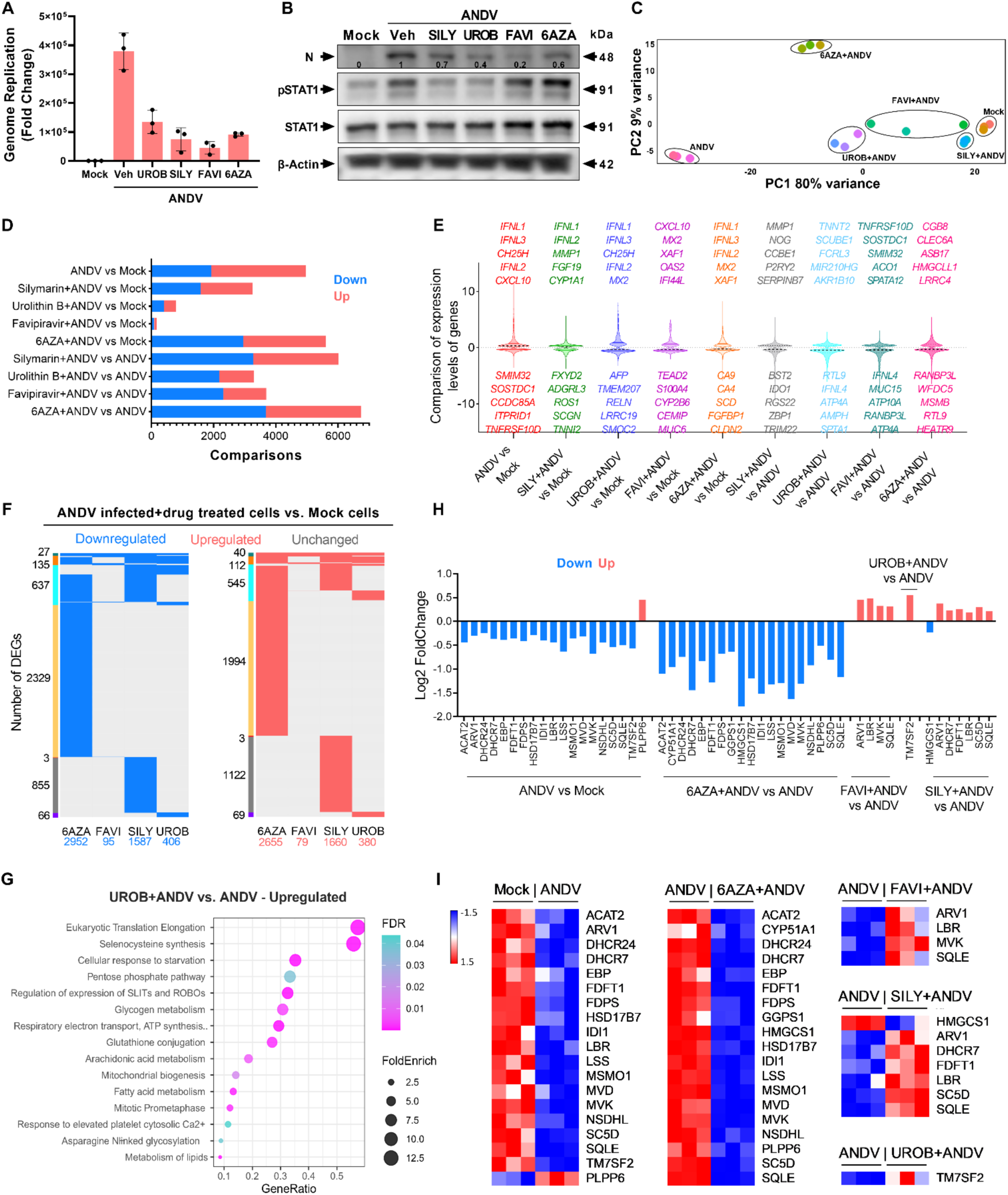
Pharmacogenomics Analysis of Compounds Exhibiting Anti-ANDV Activity in Lung Cells. **A)** The graph shows genome replication levels in indicated drug-treated ANDV-infected Calu-3 cells at 48 hpi. Dosages: UROB and SILY = 30uM, FAVI = 10uM, 6AZA = 5uM. **B)** Western blot analysis of viral N protein levels in drug-treated infected cells. Levels of host IFN pathway transactivators phospho-STAT1 and STAT1 were also detected. **C)** Principal-component analysis (PCA) plot shows global transcriptional response by drug-treated infected cells. **D)** The bar graph shows the total number of down- and up-regulated genes in indicated samples. **E)** Violin plot shows comparison of expression levels (log2 Fold Change) of DEGs in various samples. **F)** Comparison of down- and up-regulated DEGs in drug-treated infected cells relative to uninfected cells. The side bars show DEGs group (green: DEGs shared by four datasets; orange: 3 datasets; turquoise: two datasets; others: exclusive to each dataset). **G)** Dot plot shows the up-regulated pathways in urolithin B-treated infected cells relative to ANDV infected cells. **H)** Bidirectional bar chart shows DEGs involved in cholesterol biosynthesis pathway among indicated drug-treated infected cells. **I)** Heatmaps show Z scores as expression levels of DEGs involved in cholesterol biosynthesis pathway in uninfected (mock) and infected, as well as drug-treated infected, cells. Blue and red colors represent downregulated and upregulated genes, respectively. UROB = Urolithin B; SILY = Silymarin; FAVI = Favipiravir; 6AZA = 6-Azauridine.

Importantly, ANDV infection downregulated many cell cycle and metabolic pathways, specifically lipid and cholesterol biosynthesis (Figure 3A). Except 6AZA, the other three drugs were balanced or normalized with regards to the expression of several genes involved in cholesterol synthesis pathway (Figures 4H-4I, Figure S4E), suggesting that these drug-treated cells have retained cholesterol content for regular function. Expression levels of innate immune genes (IDO1, HELZ2) and inflammatory genes (IL6, ICAM1) were normalized upon drug treatment (Figure S4E). Taken together, our comprehensive pharmacogenomic analysis revealed distinct mechanisms of action for urolithin B, favipiravir, silymarin and 6AZA and pathophysiological molecular changes during ANDV infection.

## DISCUSSION

The objective of this study was to establish effective hPSC-derived modelling systems to evaluate the differences in cell tropism between Old and New World hantavirus strains. Importantly, we found that the New World hantavirus ANDV readily infected a broad variety of cell types assessed – Calu-3 lung cells, hPSC-CMs, and hPSC-astrocytes. In contrast, we found that SNV, another New World hantavirus, showed low-grade infection in all evaluated cell types. In hPSC-astrocytes and -CMs, while we did not observe much activation of innate immune pathways, pathways related to the cell cycle were readily upregulated in hPSC-CMs during SNV infection. Like SNV, the Old World HTNV showed low levels of infection in Calu-3 cells. However, both SNV and HTNV induced the activation of IFN antiviral pathways in these lung cells, which is likely a cell-type specific response to prevent more widespread infection. This promotion of IFN response has been substantiated in other independent studies^41, 42^.

The widespread and ready infection observed with ANDV across these various cell types suggests an ability by the virus to suppress antiviral response more efficiently. It has been posited that the ANDV N protein may restrict the induction of IFN-β and downstream interferon-stimulated genes (ISGs)^43^. This delay in IFN response may be the reason for rapid infection and more severe symptoms being observed in ANDV infections.

In our transcriptomic and lipidomics studies, we found that ANDV-infected cells had reduced levels of cholesterol biosynthesis genes, as well as cholesterol and triglycerides metabolites. In a previous siRNA screen of ANDV, it was found that the sterol regulatory pathway is necessary for viral entry^44^. Additional studies have confirmed that cholesterol is required for cell entry by both ANDV^45^ and HTNV^46^ and that depletion of cellular cholesterol inhibited ANDV infection^47^. One study identified a specific need for high levels of cholesterol in the pathogenesis of both ANDV and HTNV^45^. Our observation of reduction in cholesterol levels is thus exceedingly rare, indicating that additional investigation is needed to better understand the viral interaction of ANDV with the cholesterol pathway. The observed downregulation may be a host antiviral response to remove the cholesterol needed for viral replication. Other pandemic-causing and -potential viruses, including SARS-CoV-2^48^ and several types of flaviviruses^49–51^, have also been observed to require high levels of cholesterol biosynthesis for viral replication and infection. Our discovery of elevated LPC and LPE levels is consistent with similar phenomena observed over the course of SARS^52^, SARS-CoV-2^53^, and Dengue virus^54^ infection, with higher LPC and LPE levels linked to increased viral replication as a result of drastic alterations of the curvature and permeability of the cell membrane. The downregulation of cellular cholesterol and viral-mediated depletion of cholesterol we observed may explain disease outcomes in the virus. Depletion of cellular cholesterol can lead to increased cell membrane fragility, thus increasing the possibility of cell death and leakage and potentially leading to increased inflammation. Taken together with our observation of ready PSC-cardiomyocyte infection, this dysregulation of cholesterol suggests that people with cardiovascular comorbidities may be at an elevated risk of more severe infection.

Our study revealed that STING agonists, nucleoside analogues and plant-derived compounds exhibited antiviral activity. STING agonists are known to activate type 1 IFN antiviral response^53, 74^. Silymarin and urolithin B likely promote antiviral response by boosting cellular metabolism and host antiviral pathways. Silymarin antiviral activity has been demonstrated against the hepatitis C, Dengue, Chikungunya, and HIV viruses^57^. These host-directed antivirals can provide broad-spectrum protection against multiple families of viruses in the event of outbreaks. The nucleoside analog favipiravir activity against HTNV has been described previously^75^ and we confirmed its therapeutic potential against ANDV. Favipiravir and urolithin B have shown minimal transcriptomic changes upon treatment, which augurs well for their further development. Favipiravir can directly bind to hantaviral RNA-dependent RNA polymerase (RdRp) coded by the L gene. This compound also has been approved for treatment of influenza virus in Japan^76^. Thus, favipiravir inhibitory activity on RdRp is shared in multiple RNA viral families and a detailed Structure Activity Relationships (SAR) study can yield additional information on its mode of action. Similarly, direct activity of urolithin B on viral targets, if any, needs to be investigated. Despite exhibiting potent antiviral activity at non-cytotoxic doses, 6-AZA caused the largest changes in the transcriptional profile of treated cells. These unnecessary cellular changes can contribute to potential side effects. Thus, our pharmacogenomic analysis provided key safety details to make informed decisions to advance compounds to the next phases of pre-clinical and clinical testing. Additional animal studies using Syrian hamsters are needed to characterize the *in vivo* antiviral activity of these promising drug candidates.

With the rapid evolution of RNA viruses observed in the past few years, it is becoming increasingly apparent that a strong pandemic preparedness infrastructure is essential. A key tenet of this preparedness is identifying these pandemic potential viruses and those most at risk. We expect that, in the event of an ANDV pandemic, those with lung and heart comorbidities will be at an increased risk of mortality due to the broad tropism of the virus. In this study, we have generated complex *in vitro* hPSC-derived model systems and transcriptomics data, important resources which can be utilized by the research community to further advance our collective understanding of ANDV and hantaviral cell tropisms and infection mechanisms, as well as drug screening and additional compounds that can be further developed and potentially utilized in the event of an ANDV pandemic.

### Study Limitations

This study does have some limitations. First, the viral strains used were isolated from rodent hosts. Clinical isolates from humans would provide additional insights into host responses and viral replication. Obtaining these isolates from infected individuals has been difficult, however, due to infrequent infection and/or timing of diagnosis either after viral clearance or mortality^55^. Developing reverse genetics systems can help synthesize or resurrect hantaviruses based on available complete viral sequences from human infections. Second, the data presented is primarily from *in vitro* cell culture systems due to the BSL-4 containment being required to conduct animal studies using these viruses. However, we have used more advanced and biologically relevant human and animal 3D organoid systems, which can serve as valuable tools in drug development.

## Acknowledgments

We are grateful to Barbara Dillon, UCLA High Containment Program Director, for BSL-3 work. We thank Yijie Wang from the UCLA Cardiomyocyte Core for providing hPSC-CMs. We also extend our gratitude to Dr. Heinrich Feldmann from the NIH/NIAID for providing the ANDV and SNV stocks. HTNV stock was obtained from BEI Resources. We also acknowledge the CNSI Advanced Light Microscopy and Spectroscopy Lab for their support in capturing the confocal images. This study is partly supported by National Institutes of Health awards 1R01EY032149-01, 5R01AI163216-02 and 1R01DK132735-01 to VA.

## Author contributions

AJ, JIJ, SD: Collection and/or assembly of data. Data analysis and interpretation, and Manuscript writing.

NC, BK, BS, AS, CW, QC, KJW, SS, YS, AD, RD, BRS: Experimental design, Conducted experiments, Data analysis, interpretation, and Manuscript writing.

AR: Conception and design, Bioinformatics data analysis and interpretation, and Manuscript writing.

VA: Conception and design, Data analysis and interpretation, Manuscript writing and Final approval of manuscript.

## Data and materials availability

All mentioned and relevant data regarding this study is available from the above listed authors. In addition, supplementary information is available for this paper. Correspondence and requests for data and materials should be addressed to lead contact, Vaithilingaraja Arumugaswami.

## METHODS AND MATERIALS

### Ethics Statement

This study was performed in strict accordance with the recommendations of UCLA. All ANDV, SNV and HTNV live virus experiments were performed at the UCLA BSL-3 High Containment facility. Human lung tissue was obtained from deceased tissue donors in compliance with consent procedures developed by the International Institute for the Advancement of Medicine and approved by the Cedars-Sinai Medical Center Internal Review Board. The pluripotent stem cell studies were approved by the UCLA and City of Hope IRB Stem Cell Oversight Committees.

### Cells

Human lung adenocarcinoma epithelial cells (Calu-3) were purchased from the ATCC (ATCCHTB-55). These were cultured in Eagle’s Minimum Essential Medium (EMEM) (Corning), supplemented with 20 % fetal bovine serum (FBS),1 % MEM Non-Essential Amino Acids Solution (MEM NEAA), 1 % L-glutamine (L-glu) and 1 % penicillin/streptomycin (P/S). These Calu-3 cells were incubated at 37°C with 5% CO_2_. Human induced pluripotent stem cell (hiPSC) differentiation into astrocytes was induced using 0.1 μM retinoic acid (RA), 4 μM CHIR99021, 3 μM SB431542, and 2 μM Dorsomorphin for 2 days, followed by continued induction for 5 days with the removal of Dorsomorphin^56, 57^. Neural progenitor cells (NPCs) were derived from hiPSCs and further differentiated by treating them with 10as removed. The induction was continued for 5 days and NPCs were derived from hiPSCs by using 10 μM RA and SAG for 5 days. The NPC spheres were dissociated into single cells with accuse, then plated into Matrigel-coated plates at 1×10^5^ cells per well in 6-well plates. Cells were cultured for 10 days in N2B27 medium (DMEM/F12, 1xN2, 1xB27, 1xNEAA, 1xGlutamax) plus 0.1 μM RA and 1 μM SAG. They were then switched to PDGF medium (1X N2, 1XB27, 10 ng/ml PDGFAA, 5 ng/ml HGF, 10 ng/ml IGF-1, 10 ng/ml NT3, 100 ng/ml Biotin, 60 ng/ml T3, 1 μM cAMP and 25 μg/ml insulin) for another 20 days. To mature the astrocytes, the cells were switched into astrocyte maturation medium (DMEM/F12, 1xN2, 1xB27, 1xNEAA, 2 mM GlutaMAX, and10 ng/ml CNTF) for another 7 days. Human pluripotent stem cell-derived cardiomyocytes (hPSC-CMs) were provided by the UCLA Cardiomyocyte Core and were derived as described below. The hPSC-CMs were differentiated from human embryonic stem cell (hESC) line H9. The hPSCs were maintained in mTeSR1 (STEMCELL Technology) and RPMI1640 [supplemented with B27 minus insulin (Invitrogen)] was used as differentiation medium. From Days 0–1, 6 μM CHIR99021 was added into differentiation medium. On Days 3–5, 5 μM IWR1 (Sigma-Aldrich) was added to the differentiation medium. Thereafter, on Day 7, RPMI 1640 plus B27 maintenance medium was added. Finally, on Days 10–11, RPMI 1640 without D-glucose and supplemented with B27 was transiently used for metabolic purification of CMs. Distal lung epithelial organoids were cultured as described previously^35, 58^.

### Viruses

ANDV and SNV were obtained from Dr. Heinrich Feldmann from the NIH/NIAID and HTNV was obtained from BEI Resources (Table S5). ANDV, SNV, HTNV was passaged once in Vero E6 cells. Then, sequence-verified viral stocks were aliquoted and stored at -80°C. The virus titer was measured in Vero E6 cells with the established TCID50 assay.

### Viral Infection

Human lung epithelial cell line (Calu-3), hPSC-astrocytes, and hPSC-CMs were plated at 1×10^5^ cells per well using a 48-well plate. Viral inoculum of ANDV (Chile-9717869 strain), HTNV (Fojnica strain), and SNV (SNV-77734), was added onto the cells at a multiplicity of infection (MOI) of 0.1, or 1 using serum-free base media. After 1 hour incubation at 37°C with 5% CO_2_, the inoculum was replaced with respective cell type media. Cells were then fixed at selected timepoints with 4% PFA. Also, samples were collected, and RIPA buffer was used for protein analysis and Trizol for RNA isolation. For drug studies, indicated drugs were added in Calu-3 cells 24 hours prior to ANDV infection. At 48hpi, protein and RNA samples were collected for western blot and RT-qPCR analysis.

### *Viral Titer by TCID50* (Median Tissue Culture Infectious Dose) *assay*

Viral production by infected cells was quantified by the TCID50 assay, as previously described with modifications^48^. Vero E6 cells (density of 5 x10^3^cells/well) were plated in 96-well plates. The next day, viral culture media were serially diluted 10-fold (10^1^ to 10^8^) and added onto Vero E6 cells. These cells were incubated at 37°C with 5% CO_2_. 3 to 4 days after, the cells were fixed with 4% PFA and subjected to immunostaining with hantavirus anti-N antibody. The wells positive for viral infection were identified for each dilution. Then, the dilutions immediately above and below 50% of viral inhibition were determined. TCID50 was calculated based on the method of Reed and Muench.

### Drug compounds

The compounds tested were obtained from InvivoGen, Millipore Sigma and Selleckchem (Table S5). All compounds were provided as lyophilized and were then reconstituted in Nuclease-Free water (Invitrogen) or DMSO. Compounds were then aliquoted and stored at either -80°C or room temperature in dry conditions.

### Immunohistochemistry

Cells were fixed with methanol (incubated in -20°C freezer until washed with PBS) or 4% paraformaldehyde for 30-60 minutes. The cells were washed 3 times with 1x PBS and permeabilized by incubating in blocking buffer (0.3% Triton X-100, 2% BSA, 5% Goat Serum, 5% Donkey Serum in 1 X PBS) for 1 hour at room temperature. For immunostaining, cells were incubated overnight at 4°C with each primary antibody, then washed with 1X PBS three times and incubated with respective secondary antibody for 1 hour at room temperature. The cell nuclei were stained with DAPI (4’,6-Diamidino-2-Phenylindole, Dihydrochloride) (Life Technologies) at a dilution of 1:5000 in 1X PBS. Image acquisition was done using Leica DM IRB fluorescent microscopes.

### Image Analysis/Quantification

Microscope two-dimensional images were obtained using the Leica DM IL LED Fluo and Leica LAS X Software Program. 2-3 two-dimensional images were captured per well at 48hpi for each condition. These images were quantified using Image J’s plugin (Multipoint and Cell Counter). The positively stained cells were counted by a double blinded approach. Confocal slide samples were imaged using a Leica SP8 MP-DIVE-FLIM Microscope at the Advanced Light Microscopy/Spectroscopy Laboratory and Leica Microsystems Center of Excellence at the California NanoSystems Institute at UCLA (RRID:SCR_022789) with funding support from NIH Shared Instrumentation Grant S10OD025017 and NSF Major Research Instrumentation grant CHE-0722519. Confocal three-dimensional images were collected in 1024×1024 format using a 63x oil immersion objective lens, fixed scan rate of 8000Hz, and averaged 12 times. Excitation laser lines and emission detection wavelengths were optimized for the fluorescent tags as follows: blue channel excitation of 405nm with emission detection range of 420-470nm, green channel excitation of 488nm with emission detection range of 500nm-530nm, and red channel excitation of 552nm with emission detection range 590-650nm.

### Lipidomics

Cells were transferred to extraction tubes containing phosphate-buffered saline (PBS). Subsequently, a modified Bligh and Dyer extraction method (Hsieh, 2020) was employed to process the samples. Prior to the biphasic extraction step, a mixture of 70 lipid standards across 17 subclasses (AB Sciex 5040156, Avanti 330827, Avanti 330830, Avanti 330828, Avanti 791642) was added to each sample as an internal standard. Following two consecutive extractions, the pooled organic layers were dried down in Thermo SpeedVac SPD300DDA using ramp setting 4 at 35 degrees C for 45 minutes with a total run time of 90 minutes. The dried lipid samples were then resuspended in a solution of 1:1 methanol/dichloromethane with 10 mM ammonium acetate and transferred to robovials (Thermo 10800107) for subsequent analysis. The analysis of the samples was performed by direct infusion on a Sciex 5500 instrument equipped with a Differential Mobility Device (DMS)(comparable to Sciex Lipidyzer platform). A targeted acquisition list consisting of 1450 lipid species across 17 subclasses was used. The DMS was tuned with EquiSPLASH LIPIDOMIX (Avanti 330731). Data analysis was performed with in-house data analysis workflow. Detailed information regarding instrument settings, multiple reaction monitoring (MRM) lists, and analysis method are available^59, 60^. Quantitative values were normalized to cell counts.

### Phylogeny

For the phylogenetic analysis, all 68 M segment of viral sequences from *Hantaviridae, Phenuiviridae, Nairoviridae, Arenaviridae*, and *Peribunyaviridae* families of the *Bunyavirales* order (Table S1), were aligned using MAFFT v.7.505^61^ and, subsequently, these aligned sequences were used to identify GTR+F+R6 as a best-fit model based on the Bayesian Information Criteria using ModelFinder^62^. The phylogenetic tree was constructed using the maximum-likelihood (ML) method with 1,000 bootstrap replicates in IQ-TREE multi core version 2.0.3^63^. The phylogenetic tree was annotated in Interactive Tree Of Life (iTOL)^64^.

### RNA sample preparation for RNA sequencing analysis

Bulk RNA from various cell types infected with viruses, as well as drug-treated infected cells, was extracted using RNA Mini Kit (BioRad), as per the manufacturer’s instructions. RNA was quantified using a NanoDrop 2000 Spectrophotometer (Thermo Fisher Scientific). For every treatment condition, duplicate (quadruplicate samples pooled separately as duplicates) or triplicate RNA samples were submitted to the UCLA Technology Center for Genomics & Bioinformatics (TCGB) for RNA sequencing analysis.

### RNA sequencing data analysis

Library preparation, sequencing and RNA-Seq data analysis were performed as described^65, 66^ with minor modifications. In summary, libraries were prepared with the KAPA Stranded mRNA-Seq Kit, followed by second strand synthesis converting the cDNA:RNA hybrid to double-stranded cDNA (dscDNA), and incorporating dUTP into the second cDNA strand. cDNA generation was followed by end repair to generate blunt ends, A-tailing, adaptor ligation and PCR amplification. Different adaptors were used for multiplexing samples in one lane. Sequencing was performed on Illumina Novaseq 6000 for a paired-end 2×50 bp run. Data quality checking was done on Illumina SAV. Demultiplexing was performed with Illumina Bcl2fastq v2.19.1.403 software. Partek Flow ^49^ was used for all data analysis. Illumina reads from all samples were aligned to human GRCh38 reference genome using STAR 2.7.9a^50, 51^ and Ensemble transcripts release GRCh38.107 GTF was used for gene feature annotation. Subsequently, the read counts per gene were qualified.

The differential gene expression analysis was performed using DESeq2 v1.40.1 in R v4.3.0^52^ Median of ratios method was used to normalize expression counts for each gene in all samples studied. DEGs were considered if they were supported by a false discovery rate (FDR) *p* < 0.01 or pad 0.01 & FC 1. Unsupervised principal component analysis (PCA) was performed using DESeq2 in R v4.3.0. The gene ontology (GO) enrichment overrepresentation test (Release 20221013) was performed in PANTHER v17.0^53^ using PANTHER GO-SLIM Biological Process annotation dataset^54^. Reactome pathway analysis was performed using human all genes as the reference dataset in the Reactome v84^55^. GO and Reactome pathway were only considered if they were supported by FDR P < 0.05. The ggplot2 v3.4.2 in R and Prism GraphPad v9.5.1 were used to generate figures. The heatmaps were generated using pheatmap v1.0.12 in R. We deposited RNA-seq data to the NCBI GEO under the accession number GSE232641.

### Western *Blot analysis*

For protein analysis, cells were lysed in 50 mM Tris pH 7.4, 1% NP-40, 0.25% sodium deoxycholate, 1 mM EDTA, 150 mM NaCl, 1 mM Na3VO4, 20 Mm or NaF, 1mM PMSF, 2 mg ml^-1^ aprotinin, 2 mg ml^-1^ leupeptin and 0.7 mg ml^-1^ pepstatin or Laemmli Sample Buffer (Bio Rad, Hercules, CA). Cell lysates were resolved by SDS-PAGE using 10% gradient gels (Bio-Rad), then transferred to a 0.2 µm PVDF membrane (Bio-Rad). After the transfer, the membranes were blocked (5% skim milk and 0.1% Tween-20) in 1x TBST (0.1% Tween-20) at room temperature (RT) for 1 hour. The membranes were then incubated with the respective monoclonal antibodies overnight at 4°C and detected by SuperSignal West Femto Maximum Sensitivity Substrate (Thermo Scientific). Membranes were exposed and visualized with the Bio-Rad ChemiDoc MP Imaging System.

### Statistics and Data analysis

GraphPad Prism, version 9.5.1 was used for graph generation and statistical analysis. Data was then analyzed for statistical significance using an unpaired student’s *t*-test to compare two groups (uninfected vs. infected) or a non-parametric t-test (Mann-Whitney Test). All statistical testing were performed at the two-sided alpha level of 0.05.

## REFERENCES

1. Avšič-Županc T, Saksida A, Korva M. Hantavirus infections. Clinical Microbiology and Infection. 2019;21:e6–e16. doi:10.1111/1469-0691.12291

2. Watson DC, Sargianou M, Papa A, Chra P, Starakis I, Panos G. Epidemiology of Hantavirus infections in humans: A comprehensive, global overview. Critical Reviews in Microbiology. 2014;40(3):261–272. doi:10.3109/1040841X.2013.783555

3. Mir M. Hantaviruses. Clin Lab Med. 2010;30(1):67-91. doi:10.1016/j.cll.2010.01.004

4. Lee HW, Lee PW, Johnson KM. Isolation of the Etiologic Agent of Korean Hemorrhagic Fever. The Journal of Infectious Diseases. 1978;137(3):298–308. doi:10.1093/infdis/137.3.298

5. Nichol ST, Spiropoulou CF, Morzunov S, et al. Genetic Identification of a Hantavirus Associated with an Outbreak of Acute Respiratory Illness. Science. 1993;262(5135):914-917. doi:10.1126/science.8235615

6. Nerurkar VivekR, Song KJ, Carleton Gajdusek D, Yanagihara R. Genetically distinct hantavirus in deer mice. The Lancet. 1993;342(8878):1058–1059. doi:10.1016/0140-6736(93)92917-I

7. Brummer-Korvenkontio M, Vaheri A, Hovi T, et al. Nephropathia Epidemica: Detection of Antigen in Bank Voles and Serologic Diagnosis of Human Infection. The Journal of Infectious Diseases. 1980;141(2):131–134. doi:10.1093/infdis/141.2.131

8. Calderón G, Pini N, Bolpe J, et al. Hantavirus Reservoir Hosts Associated with Peridomestic Habitats in Argentina - Volume 5, Number 6—December 1999 - Emerging Infectious Diseases journal - CDC. doi:10.3201/eid0506.990608

9. Jonsson CB, Figueiredo LTM, Vapalahti O. A Global Perspective on Hantavirus Ecology, Epidemiology, and Disease. Clin Microbiol Rev. 2010;23(2):412–441. doi:10.1128/CMR.00062-09

10. Enría D, Padula P, Segura EL, et al. Hantavirus pulmonary syndrome in Argentina. Possibility of person to person transmission. Medicina (B Aires). 1996;56(6):709–711.

11. Wells RM, Sosa Estani S, Yadon ZE, et al. An unusual hantavirus outbreak in southern Argentina: person-to-person transmission? Hantavirus Pulmonary Syndrome Study Group for Patagonia. Emerg Infect Dis. 1997;3(2):171–174. doi:10.3201/eid0302.970210

12. Elliott RM. Molecular biology of the Bunyaviridae. J Gen Virol. 1990;71 (Pt 3):501–522. doi:10.1099/0022-1317-71-3-501

13. Dieterle ME, Solà-Riera C, Ye C, et al. Genetic depletion studies inform receptor usage by virulent hantaviruses in human endothelial cells. Elife. 2021;10:e69708. doi:10.7554/eLife.69708

14. Gavrilovskaya IN, Shepley M, Shaw R, Ginsberg MH, Mackow ER. beta3 Integrins mediate the cellular entry of hantaviruses that cause respiratory failure. Proc Natl Acad Sci U S A. 1998;95(12):7074–7079. doi:10.1073/pnas.95.12.7074

15. Jangra RK, Herbert AS, Li R, et al. Protocadherin-1 is essential for cell entry by New World hantaviruses. Nature. 2018;563(7732):559–563. doi:10.1038/s41586-018-0702-1

16. Noack D, Goeijenbier M, Reusken CBEM, Koopmans MPG, Rockx BHG. Orthohantavirus Pathogenesis and Cell Tropism. Front Cell Infect Microbiol. 2020;10:399. doi:10.3389/fcimb.2020.00399

17. Zaki SR, Greer P w., Coffield LM, et al. Hantavirus Pulmonary Syndrome. Am J Pathol. 1995;146(3):552–579.

18. Green W, Feddersen R, Yousef O, et al. Tissue distribution of hantavirus antigen in naturally infected humans and deer mice. J Infect Dis. 1998;177(6):1696–1700. doi:10.1086/515325

19. Toro J, Vega JD, Khan AS, et al. An outbreak of hantavirus pulmonary syndrome, Chile, 1997. Emerg Infect Dis. 1998;4(4):687–694.

20. Nolte KB, Feddersen RM, Foucar K, et al. Hantavirus pulmonary syndrome in the United States: a pathological description of a disease caused by a new agent. Hum Pathol. 1995;26(1):110–120. doi:10.1016/0046-8177(95)90123-x

21. Saggioro FP, Rossi MA, Duarte MIS, et al. Hantavirus infection induces a typical myocarditis that may be responsible for myocardial depression and shock in hantavirus pulmonary syndrome. J Infect Dis. 2007;195(10):1541–1549. doi:10.1086/513874

22. Easterbrook JD, Klein SL. Seoul virus enhances regulatory and reduces proinflammatory responses in male Norway rats. J Med Virol. 2008;80(7):1308–1318. doi:10.1002/jmv.21213

23. Yanagihara R, Amyx HL, Gajdusek DC. Experimental infection with Puumala virus, the etiologic agent of nephropathia epidemica, in bank voles (Clethrionomys glareolus). J Virol. 1985;55(1):34–38.

24. Maas M, van Heteren M, de Vries A, et al. Seoul Virus Tropism and Pathology in Naturally Infected Feeder Rats. Viruses. 2019;11(6):E531. doi:10.3390/v11060531

25. Mackow ER, Gavrilovskaya IN. Hantavirus regulation of endothelial cell functions. Thromb Haemost. 2009;102(6):1030–1041. doi:10.1160/TH09-09-0640

26. Spiropoulou CF, Srikiatkhachorn A. The role of endothelial activation in dengue hemorrhagic fever and hantavirus pulmonary syndrome. Virulence. 2013;4(6):525–536. doi:10.4161/viru.25569

27. Fosse JH, Haraldsen G, Falk K, Edelmann R. Endothelial Cells in Emerging Viral Infections. Front Cardiovasc Med. 2021;8:619690. doi:10.3389/fcvm.2021.619690

28. Riquelme R. Hantavirus. Semin Respir Crit Care Med. 2021;42(6):822–827. doi:10.1055/s-0041-1733803

29. Hautala T, Partanen T, Sironen T, et al. Elevated cerebrospinal fluid neopterin concentration is associated with disease severity in acute Puumala hantavirus infection. Clin Dev Immunol. 2013;2013:634632. doi:10.1155/2013/634632

30. Partanen T, Chen J, Lehtonen J, et al. Heterozygous TLR3 Mutation in Patients with Hantavirus Encephalitis. J Clin Immunol. 2020;40(8):1156–1162. doi:10.1007/s10875-020-00834-2

31. Gohil J, Gowda A, George T, Easwer HV, George A, Nair P. Pituitary apoplexy and panhypopituitarism following acute leptospirosis. Pituitary. 2021;24(6):854–858. doi:10.1007/s11102-021-01156-1

32. Talamonti L, Padula PJ, Canteli MS, Posner F, Marczeski FP, Weller C. Hantavirus pulmonary syndrome: encephalitis caused by virus Andes. J Neurovirol. 2011;17(2):189–192. doi:10.1007/s13365-010-0011-4

33. Shin OS, Song GS, Kumar M, Yanagihara R, Lee HW, Song JW. Hantaviruses Induce Antiviral and Pro-Inflammatory Innate Immune Responses in Astrocytic Cells and the Brain. Viral Immunol. 2014;27(6):256–266. doi:10.1089/vim.2014.0019

34. Liu Y yuan, Chen L jun, Zhong Y, et al. Specific interference shRNA-expressing plasmids inhibit Hantaan virus infection in vitro and in vivo. Acta Pharmacol Sin. 2016;37(4):497–504. doi:10.1038/aps.2015.165

35. Mulay A, Konda B, Garcia G, et al. SARS-CoV-2 infection of primary human lung epithelium for COVID-19 modeling and drug discovery. Cell Reports. 2021;35(5):109055. doi:10.1016/j.celrep.2021.109055

36. Botten J, Mirowsky K, Kusewitt D, et al. Experimental infection model for Sin Nombre hantavirus in the deer mouse (Peromyscus maniculatus). Proc Natl Acad Sci U S A. 2000;97(19):10578–10583.

37. Botten J, Mirowsky K, Kusewitt D, et al. Persistent Sin Nombre Virus Infection in the Deer Mouse (Peromyscus maniculatus) Model: Sites of Replication and Strand-Specific Expression. J Virol. 2003;77(2):1540–1550. doi:10.1128/JVI.77.2.1540-1550.2002

38. Schountz T, Acuña-Retamar M, Feinstein S, et al. Kinetics of Immune Responses in Deer Mice Experimentally Infected with Sin Nombre Virus. J Virol. 2012;86(18):10015–10027. doi:10.1128/JVI.06875-11

39. McGuire A, Miedema K, Fauver JR, et al. Maporal Hantavirus Causes Mild Pathology in Deer Mice (Peromyscus maniculatus). Viruses. 2016;8(10):E286. doi:10.3390/v8100286

40. Mittler E, Dieterle ME, Kleinfelter LM, Slough MM, Chandran K, Jangra RK. Hantavirus entry: Perspectives and recent advances. Adv Virus Res. 2019;104:185–224. doi:10.1016/bs.aivir.2019.07.002

41. Brocato RL, Wahl V, Hammerbeck CD, et al. Innate immune responses elicited by Sin Nombre virus or type I IFN agonists protect hamsters from lethal Andes virus infections. J Gen Virol. Published online August 1, 2018. doi:10.1099/jgv.0.001131

42. Brocato RL, Altamura LA, Carey BD, et al. Comparison of transcriptional responses between pathogenic and nonpathogenic hantavirus infections in Syrian hamsters using NanoString. PLoS Negl Trop Dis. 2021;15(8):e0009592. doi:10.1371/journal.pntd.0009592

43. Alff PJ, Sen N, Gorbunova E, Gavrilovskaya IN, Mackow ER. The NY-1 Hantavirus Gn Cytoplasmic Tail Coprecipitates TRAF3 and Inhibits Cellular Interferon Responses by Disrupting TBK1-TRAF3 Complex Formation. J Virol. 2008;82(18):9115–9122. doi:10.1128/JVI.00290-08

44. Petersen J, Drake MJ, Bruce EA, et al. The Major Cellular Sterol Regulatory Pathway Is Required for Andes Virus Infection. PLoS Pathog. 2014;10(2):e1003911. doi:10.1371/journal.ppat.1003911

45. Kleinfelter LM, Jangra RK, Jae LT, et al. Haploid Genetic Screen Reveals a Profound and Direct Dependence on Cholesterol for Hantavirus Membrane Fusion. mBio. 2015;6(4):e00801–15. doi:10.1128/mBio.00801-15

46. Krautkrämer E, Zeier M. Hantavirus Causing Hemorrhagic Fever with Renal Syndrome Enters from the Apical Surface and Requires Decay-Accelerating Factor (DAF/CD55). J Virol. 2008;82(9):4257–4264. doi:10.1128/JVI.02210-07

47. Chiang CF, Flint M, Lin JMS, Spiropoulou CF. Endocytic Pathways Used by Andes Virus to Enter Primary Human Lung Endothelial Cells. PLoS One. 2016;11(10):e0164768. doi:10.1371/journal.pone.0164768

48. Daniloski Z, Jordan TX, Wessels HH, et al. Identification of Required Host Factors for SARS-CoV-2 Infection in Human Cells. Cell. 2021;184(1):92–105.e16. doi:10.1016/j.cell.2020.10.030

49. Martín-Acebes MA, Vázquez-Calvo Á, Saiz JC. Lipids and flaviviruses, present and future perspectives for the control of dengue, Zika, and West Nile viruses. Progress in Lipid Research. 2016;64:123–137. doi:10.1016/j.plipres.2016.09.005

50. Singh PK, Khatri I, Jha A, et al. Determination of system level alterations in host transcriptome due to Zika virus (ZIKV) Infection in retinal pigment epithelium. Sci Rep. 2018;8:11209. doi:10.1038/s41598-018-29329-2

51. Soto-Acosta R, Bautista-Carbajal P, Cervantes-Salazar M, Angel-Ambrocio AH, Del Angel RM. DENV up-regulates the HMG-CoA reductase activity through the impairment of AMPK phosphorylation: A potential antiviral target. PLoS Pathog. 2017;13(4):e1006257. doi:10.1371/journal.ppat.1006257

52. Wu Q, Zhou L, Sun X, et al. Altered Lipid Metabolism in Recovered SARS Patients Twelve Years after Infection. Sci Rep. 2017;7:9110. doi:10.1038/s41598-017-09536-z

53. Barberis E, Timo S, Amede E, et al. Large-Scale Plasma Analysis Revealed New Mechanisms and Molecules Associated with the Host Response to SARS-CoV-2. Int J Mol Sci. 2020;21(22):8623. doi:10.3390/ijms21228623

54. Wang X, Nijman R, Camuzeaux S, et al. Plasma lipid profiles discriminate bacterial from viral infection in febrile children. Sci Rep. 2019;9:17714. doi:10.1038/s41598-019-53721-1

55. Galeno H, Mora J, Villagra E, et al. First Human Isolate of Hantavirus (Andes virus) in the Americas. Emerg Infect Dis. 2002;8(7):657–661. doi:10.3201/eid0807.010277

56. Li L, Tian E, Chen X, et al. GFAP Mutations in Astrocytes Impair Oligodendrocyte Progenitor Proliferation and Myelination in an hiPSC Model of Alexander Disease. Cell Stem Cell. 2018;23(2):239–251.e6. doi:10.1016/j.stem.2018.07.009

57. Wang C, Zhang M, Garcia G, et al. ApoE-Isoform-Dependent SARS-CoV-2 Neurotropism and Cellular Response. Cell Stem Cell. 2021;28(2):331–342.e5. doi:10.1016/j.stem.2020.12.018

58. Konda B, Mulay A, Yao C, Beil S, Israely E, Stripp BR. Isolation and Enrichment of Human Lung Epithelial Progenitor Cells for Organoid Culture. J Vis Exp. 2020;(161). doi:10.3791/61541

59. Su B, Bettcher LF, Hsieh WY, et al. A DMS Shotgun Lipidomics Workflow Application to Facilitate High-Throughput, Comprehensive Lipidomics. J Am Soc Mass Spectrom. 2021;32(11):2655–2663. doi:10.1021/jasms.1c00203

60. Hsieh WY, Williams KJ, Su B, Bensinger SJ. Profiling of mouse macrophage lipidome using direct infusion shotgun mass spectrometry. STAR Protocols. 2021;2(1):100235. doi:10.1016/j.xpro.2020.100235

61. Katoh K, Rozewicki J, Yamada KD. MAFFT online service: multiple sequence alignment, interactive sequence choice and visualization. Brief Bioinform. 2019;20(4):1160–1166. doi:10.1093/bib/bbx108

62. Kalyaanamoorthy S, Minh BQ, Wong TKF, von Haeseler A, Jermiin LS. ModelFinder: fast model selection for accurate phylogenetic estimates. Nat Methods. 2017;14(6):587–589. doi:10.1038/nmeth.4285

63. Minh BQ, Schmidt HA, Chernomor O, et al. IQ-TREE 2: New Models and Efficient Methods for Phylogenetic Inference in the Genomic Era. Mol Biol Evol. 2020;37(5):1530–1534. doi:10.1093/molbev/msaa015

64. Letunic I, Bork P. Interactive Tree Of Life (iTOL) v5: an online tool for phylogenetic tree display and annotation. Nucleic Acids Res. 2021;49(W1):W293–W296. doi:10.1093/nar/gkab301

65. Garcia G, Irudayam JI, Jeyachandran AV, et al. Innate immune pathway modulator screen identifies STING pathway activation as a strategy to inhibit multiple families of arbo and respiratory viruses. Cell Rep Med. 2023;4(5):101024. doi:10.1016/j.xcrm.2023.101024

66. Wu B, Ramaiah A, Garcia G, Hasiakos S, Arumugaswami V, Srikanth S. ORAI1 Limits SARS-CoV-2 Infection by Regulating Tonic Type I IFN Signaling. J Immunol. 2022;208(1):74–84. doi:10.4049/jimmunol.2100742

