## Supplemental Materials File for "Comparative Analysis of Molecular Pathogenic Mechanisms and Antiviral Development Targeting Old and New World Hantaviruses"

### SUPPLEMENTAL FIGURES

#### Supplementary Figure 1

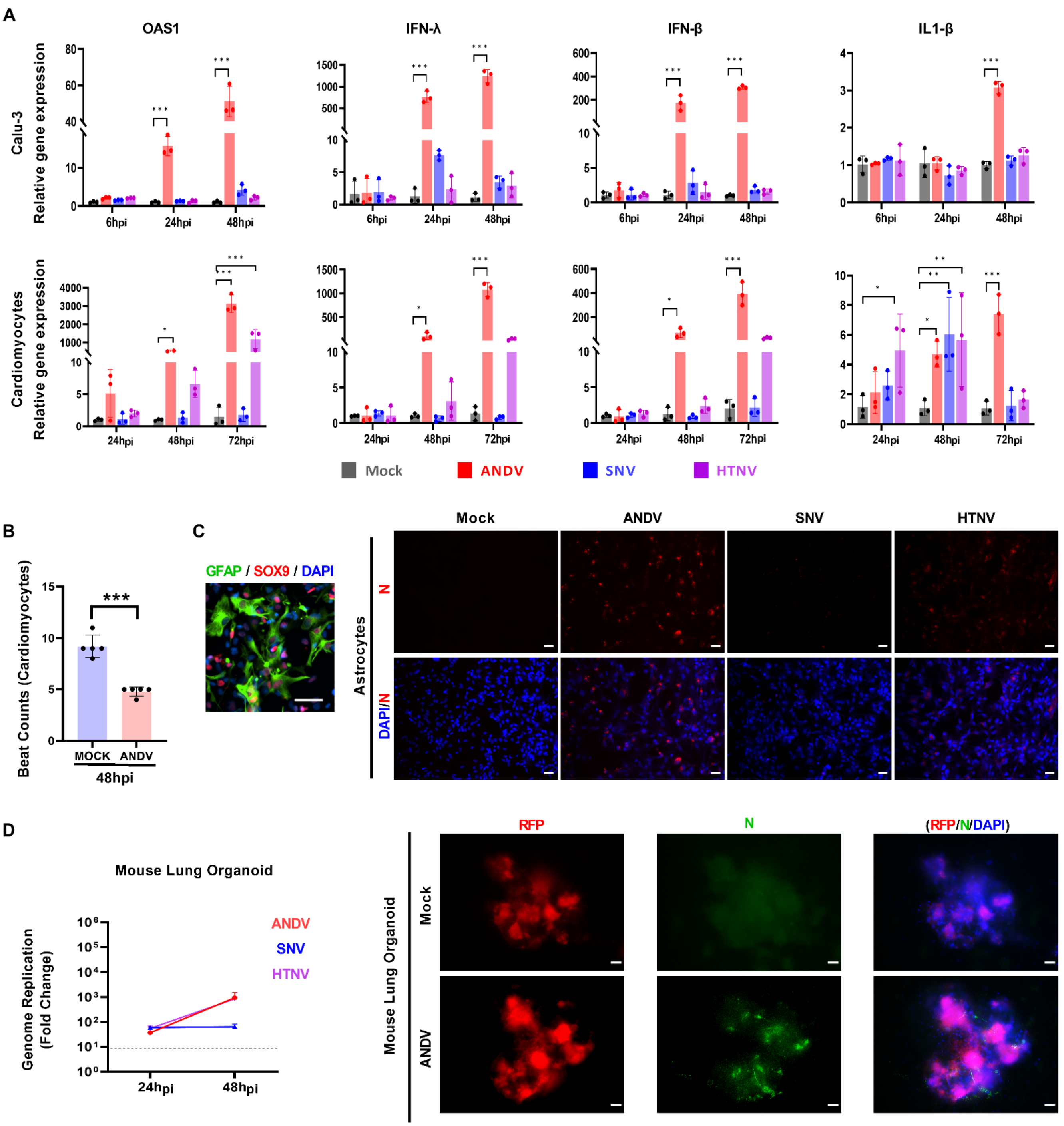


**Supplementary Fig. 1. Evaluating Cell Tropism of Various Hantaviruses. A)** Graphs represent relative expression of various immune genes in Calu-3 cells and hPSC-CMs.

Quantitative data are presented as mean ± standard deviation. Statistical comparisons were made using ANOVA followed by Tukey's post hoc test (*, $P < 0.05$; **, $P < 0.001$). **B)** The Graph shows the count of IPSC-CM beats in Mock and ANDV-infected cells in 48hpi. **C)** Immunofluorescent images of hPSC-astrocytes. (Left) Cells were confirmed with astrocyte-specific markets GFAP (green) and SOX9 (red). (Right) These cells were infected with indicated viruses at 48 hpi. Red = viral N protein. Scale bar: 25μm. **D)** The graph shows the comparative genome replication of viruses in mouse lung organoids at indicated timepoints. Fluorescence microscopy images show ANDV-infected (green = N protein) mouse lung organoids expressing RFP transgene at 48 hpi. Scale bar: 25μm. Quantitative data are presented as mean ± standard deviation. Statistical analysis was performed using ANOVA, followed by Tukey's post hoc test (*, $P < 0.05$; **, $P < 0.01$; ***, $P < 0.001$).

**Supplementary Figure 2**

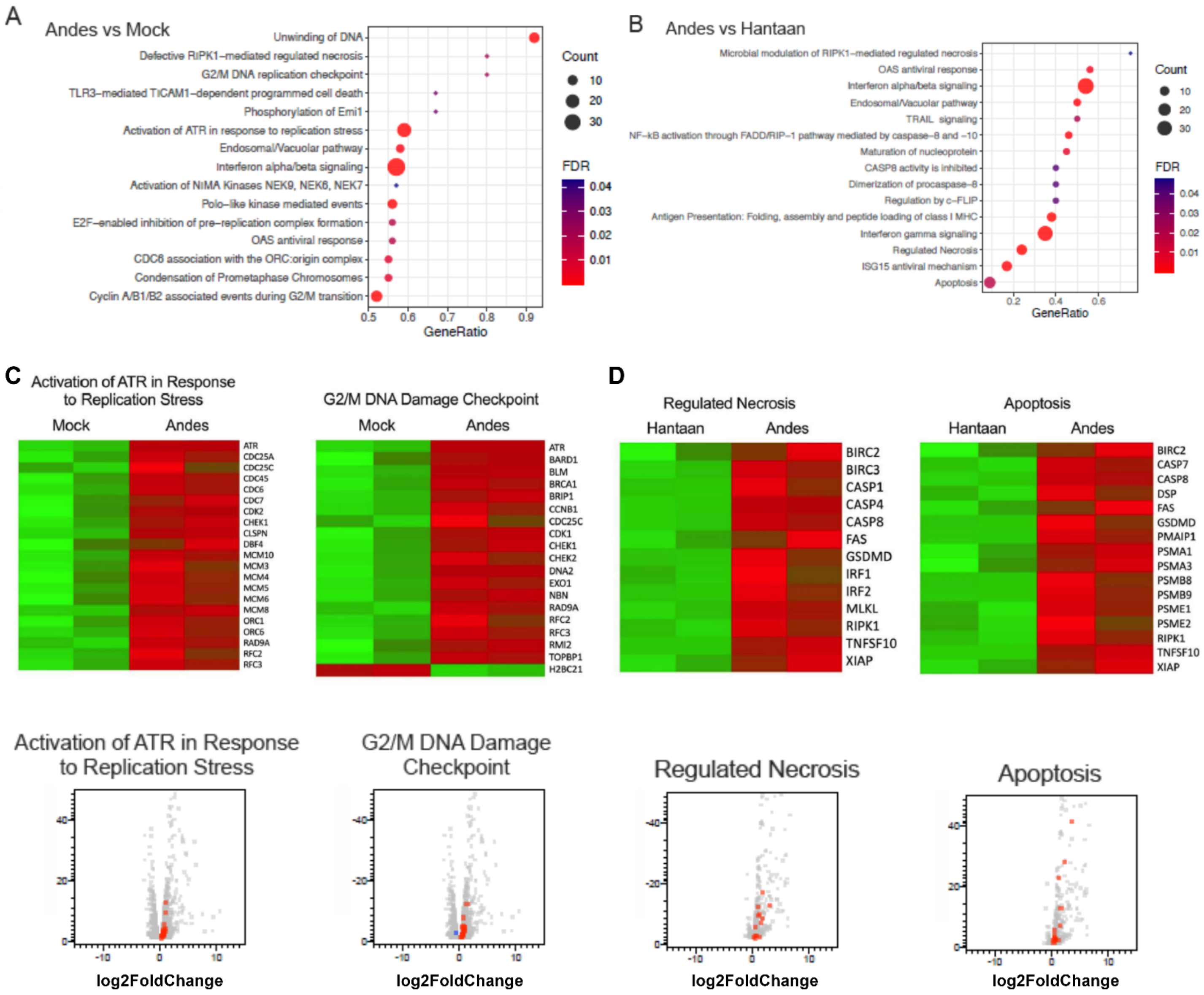


**Supplementary Fig. 2. Transcriptomic Analysis of Hantaviral Infection in hPSC-CMs. A, B)** The dot plot shows the pathway analysis of ANDV-infected/mock-infected and ANDV-infected/HTNV-infected hPSC-CMs at 48 hpi. **C, D)** Heatmap illustrates Z scores as expression levels of the genes involved in the indicated pathways in mock and infected hPSC-CMs. Red and green represent upregulated and downregulated genes, respectively. The corresponding volcano plots illustrates the differential expression of statistically significant genes of these pathways**.**

**Supplementary Figure 3**

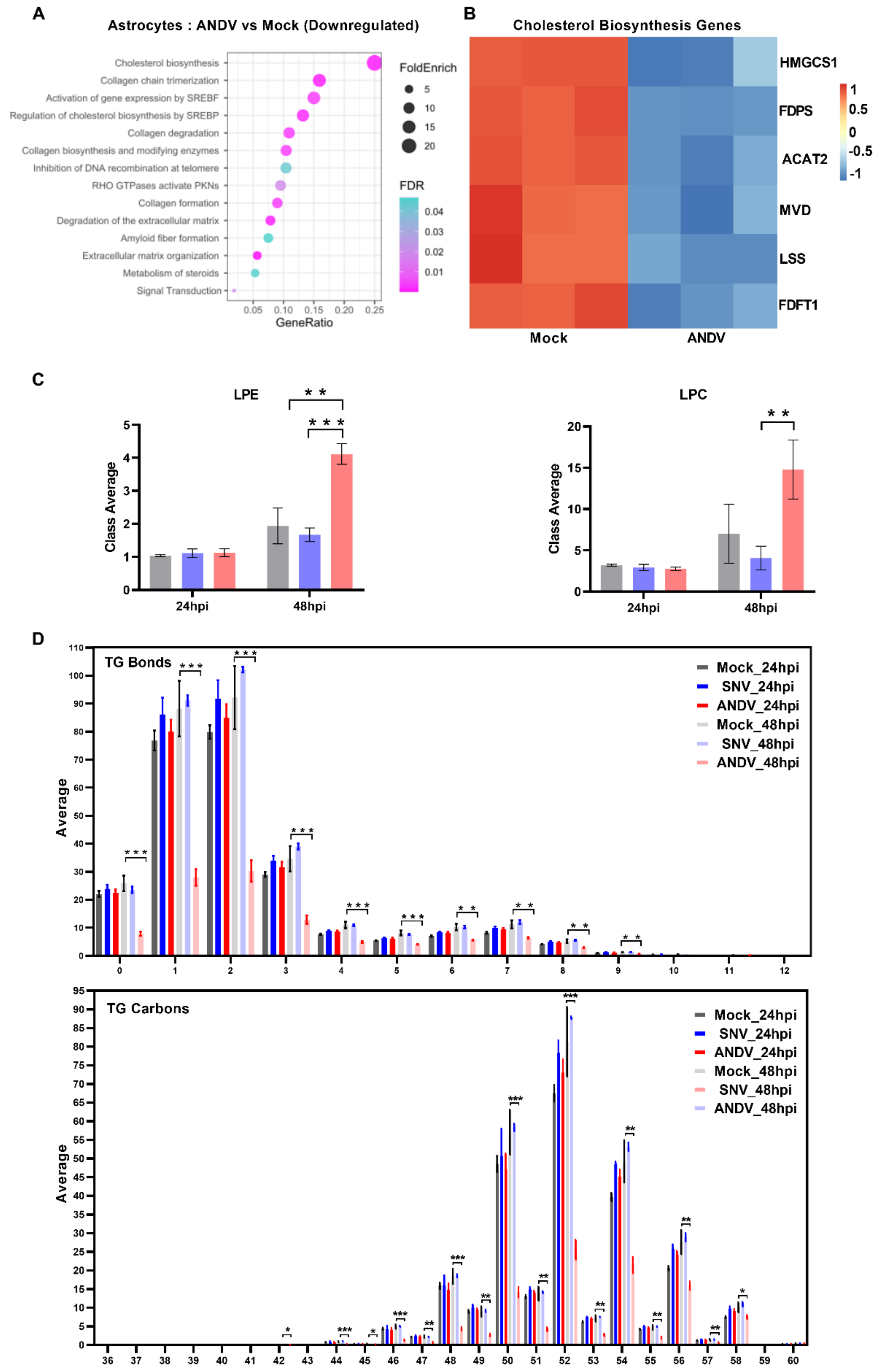


**Supplementary Fig. 3. Examining Hantavirus-Induced Dysregulation of Cholesterol Biosynthesis and Metabolism Pathways. A)** The dot plot depicts the impact of ANDV on expression of cholesterol pathway-associated genes and other indicated cellular pathways in hPSC-astrocytes. **B)** Heatmap illustrating Z scores representing reduced expression levels of genes associated with the cholesterol biosynthesis pathway. Red and blue correspond to up- and down-regulation, respectively. **C)** The graphs show the levels of lipid metabolites, LPE (Lysophosphatidylethanolamine) and LPC (Lysophosphatidylcholine), in mock as well as SNV or ANDV-infected Calu-3 cells. **D)** The graphs show the triglyceride (TG) bonds and TG carbon levels present in the mock and infected Calu-3 cells at 24 and 48 hpi. Quantitative data are presented as mean ± standard deviation. Statistical analysis was performed using ANOVA, followed by Tukey's post hoc test *, $P < 0.05$; **, $P < 0.01$; ***, $P < 0.001$).

#### Supplementary Figure 4

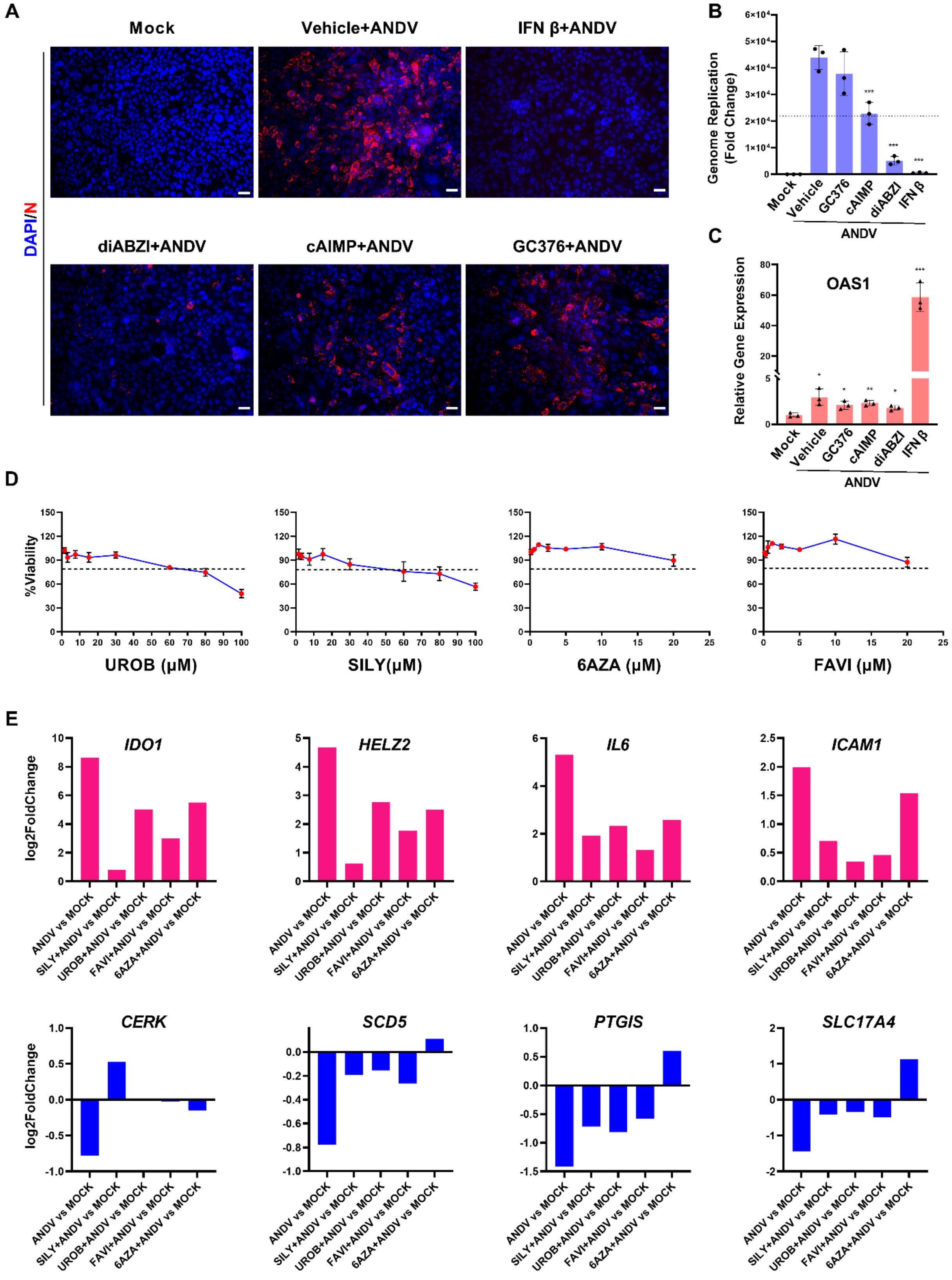

**Supplementary Fig. 4. Pharmacogenomic Analysis of Potential Anti-ANDV Drug Compounds. A)** Immunofluorescence images of ANDV-infected Calu-3 cells treated with vehicle or various drug compounds at 24 hpi. Red = N protein. Scale bar = 25μm. **B)** The graph shows the levels of viral genome replication at 24 hpi in response to indicated drug compounds. **C)** The graph represents relative gene expression levels of the innate immune gene OAS1 in response to treatment with indicated drug compounds. **D)** The graphs illustrate the dose-response viability assay of indicated drug compounds in Calu-3 cells at 48 hours post-treatment. **E)** The bar charts show the varying Log2 Fold Change values of indicated genes in ANDV-infected, as well as drug-treated infected, cells at 48hpi. Quantitative data are presented as mean ± standard deviation. Statistical analysis was performed using ANOVA, followed by Tukey's post hoc test (*, $P < 0.05$; **, $P < 0.001$).

### SUPPLEMENTARY TABLES

#### Supplementary Table 1

| VIRUS_NAME | ACCESSION_NUMBER | FAMILY | STRAIN |
|---|---|---|---|
| Necocli_virus | NC_043408.1 | Hantaviridae | HV O0020002 |
| Yakeshi_virus | NC_038705.1 | Hantaviridae | Yakeshi-Si-210 |
| Rockport_virus | NC_038694.1 | Hantaviridae | MSB57412 |
| Oxbow_virus | NC_043176.1 | Hantaviridae | Ng1453 |
| Eothenomys_miletus_hantavirus | NC_038528.1 | Hantaviridae | LX309 |
| Fusong_Hantavirus | NC_038447.1 | Hantaviridae | Mf-682 |
| Dabieshan_virus | NC_038383.1 | Hantaviridae | Yongjia-Nc-58 |
| Choclo_virus | NC_038374.1 | Hantaviridae | Not available |
| Asama_virus | NC_038274.1 | Hantaviridae | N10 |
| Amga_virus | KF974359.1 | Hantaviridae | AH301 |
| Kenkeme_virus | NC_034565.1 | Hantaviridae | Fuyuan-Sr-326 |
| Fugong_virus | NC_034466.1 | Hantaviridae | FG10 |
| Bowe_virus | NC_034406.1 | Hantaviridae | VN1512 |
| Bruges_hantavirus | NC_034395.1 | Hantaviridae | BE/Vieux-Genappe/TE/2013/1 |
| Jeju_virus | NC_034404.1 | Hantaviridae | 10-11 |
| Lena_River_virus | MH499471.2 | Hantaviridae | Not available |
| Tatenale_orthohantavirus | NC_055637.1 | Hantaviridae | Upton_Heath |
| Robina_orthohantavirus | NC_055634.1 | Hantaviridae | P17-14855 |
| Prospect_Hill_virus | NC_038940.1 | Hantaviridae | Not available |
| Laguna_Negra_virus | NC_038506.1 | Hantaviridae | Not available |
| El_Moro_Canyon_hantavirus | NC_038424.1 | Hantaviridae | RM-97 |
| Black_Creek_Canal_virus | NC_043073.1 | Hantaviridae | Not available |
| Bayou_virus | NC_038300.1 | Hantaviridae | Not available |
| Asikkala_virus | NC_043069.1 | Hantaviridae | CZ/Beskydy/412/2010/Sm |
| Dobrava_virus | NC_005234.1 | Hantaviridae | DOBV/Ano-Poroia/Afl9/1999 |
| Hantaan_virus | NC_005219.1 | Hantaviridae | Not available |
| Seoul_virus | NC_005237.1 | Hantaviridae | Not available |
| Tula_virus | NC_005228.1 | Hantaviridae | Not available |
| Sin_Nombre_virus | NC_005215.1 | Hantaviridae | Not available |
| Puumala_virus | NC_005223.1 | Hantaviridae | Not available |
| Andes_virus | NC_003467.2 | Hantaviridae | Not available |
| Anjozorobe_hantavirus | NC_034563.1 | Hantaviridae | Anjozorobe/Em/MDG/2009/ATD49 |
| Cano_Delgadito_virus | NC_034525.1 | Hantaviridae | VHV-574 |
| Khabarovsk_virus | NC_034518.1 | Hantaviridae | Fuyuan-Mm-217 |
| Sangassou_virus | NC_034516.1 | Hantaviridae | SA14 |
| Cao_Bang_virus | NC_034474.1 | Hantaviridae | Not available |
| Montano_virus | NC_034397.1 | Hantaviridae | 104/2006 |
| Andes_virus | AY228238.1 | Hantaviridae | CHI-7913 |
| Hantaanvirus | EU092222.1 | Hantaviridae | CGHu1 |

| | | | |
|---|---|---|---|
| Sin_Nombre_virus | L37903.1 | Hantaviridae | NM R11 |
| Wenling_frogfish_arenavirus | NC_040466.1 | Arenaviridae | XYHYG24857 |
| Wenling_frogfish_arenavirus | NC_040426.1 | Arenaviridae | XYHYG11303 |
| Abu_Hammad_virus | KU925435.1 | Nairoviridae | Art 1194 |
| Abu_Mina_virus | KU925438.1 | Nairoviridae | EG AN 4996 |
| Artashat_virus | NC_043442.1 | Nairoviridae | LEIV-10898Az |
| Avalon_virus | NC_040459.1 | Nairoviridae | CanAr173 |
| Bandia_virus | KU925447.1 | Nairoviridae | RV611 |
| Burana_virus | NC_043438.1 | Nairoviridae | 760 |
| Chim_virus | NC_043436.1 | Nairoviridae | LEIV-858Uz |
| Crimean_Congo_hemorrhagic_fever_virus | DQ211625.1 | Nairoviridae | AP92 |
| Dera_Ghazi_Khan_virus | NC_034510.1 | Nairoviridae | JD254 |
| Bhanja_virus | NC_027141.1 | Phenuiviridae | ibAr2709 |
| Herbert_virus | NC_038713.1 | Peribunyaviridae | F23/CI/2004 |
| Tai_virus | NC_034461.1 | Peribunyaviridae | F47/CI/2004 |
| Kibale_virus | NC_034468.1 | Peribunyaviridae | P05/UG/2008 |
| Lakamha_virus | MN092356.1 | Peribunyaviridae | Palenque-C559-MX-2008 |
| Abras_virus | MH017281.1 | Peribunyaviridae | 75V1183 |
| Bakau_virus | MK896634.1 | Peribunyaviridae | Not available |
| Telok_Forest_virus | MK896452.1 | Peribunyaviridae | Not available |
| Moju_virus | KP792674.1 | Peribunyaviridae | BeAr 12590 |
| Caimito_virus | NC_055410.1 | Peribunyaviridae | VP-488A |
| Hubei_diptera_virus | NC_032159.1 | Phenuiviridae | SCM17647 |
| Cumuto_virus | NC_043046.1 | Phenuiviridae | TR7904 |
| Hubei_diptera_virus | NC_032278.1 | Phenuiviridae | SCM94992 |
| Hubei_lepidoptera_virus | NC_032257.1 | Phenuiviridae | LCM141331 |
| Badu_virus | NC_038258.1 | Phenuiviridae | TS6347 |
| Alenquer_virus | NC_055332.1 | Phenuiviridae | Not available |
| Candiru_virus | NC_015373.1 | Phenuiviridae | Not available |

**Supplementary Table 1. *Bunyavirales* Order Viruses Used for ML Phylogeny.** Details of M gene sequences (n = 68) of various viruses belonging to the *Bunyavirales* order used from NCBI to construct the ML phylogeny (Related to Figure 1A). Each row is color-coded to match the phylogenetic cluster colors in Figure 1A.

**Supplementary Table 2. Commonly Upregulated Genes in Cell Types Infected with Various Cell Types at 48 hpi.** List of all the commonly upregulated gene counts in ANDV-, SNV- and HTNV-infected cell types at 48 hpi (Related to Figure 2). (Attached Separately).

**Supplementary Table 3. Commonly Up- and Downregulated Hantaviral Cell Entry Receptors in Infected Cell Types at 48 hpi.** List of all the commonly up- and down-regulated putative hantaviral cell entry receptor genes in ANDV-, SNV- and HTNV-infected cell types at 48 hpi. (Attached Separately).

**Supplementary Table 4. Hantaviral Annotation of DEG Sets in Each Cell Type and Drug Treatment Using PANTHER and Overrepresented Downregulated Reactome Pathways.** Hantaviral annotation of DEG sets in each cell type and drug treatment performed using PANTHER, as well as the list of overrepresented downregulated Reactome Pathways, are presented (Related to Figure 4). (Attached Separately).

**Supplementary Table 5:**

| REAGENT/RESOURCE | SOURCE | IDENTIFIER |
|---|---|---|
| **Antibodies** | | |
| Polyclonal Anti-Sin Nombre Virus, SN77734 Nucleocapsid Protein Antibody | BEI Resources | Cat#NR-9676 |
| Anti-Cardiac Troponin T antibody [1F11] | Abcam | Cat#ab10214 |
| Anti-Pro-SP-C - Rabbit, n-terminal Antibody | Seven Hills Bioreagents | Cat#WRAB-9337 |
| Cleaved caspase-3 rabbit monoclonal antibody, clone D175 | Cell Signaling Technology | Cat#9661S |
| Phospho-Stat1 (Tyr701) (58D6) Rabbit mAb | Cell Signaling | Cat#9167S |
| Stat1 (D1K9Y) Rabbit mAb | Cell Signaling | Cat#14994S |
| Sin Nombre Virus Nucleocapsid Antibody - BSA Free | Novus Biologicals | Cat# NBP2-41257 |
| Goat anti-Mouse IgG (H+L) Cross-Adsorbed Secondary Antibody, Alexa Fluor 555 | Thermo Fisher Scientific | Cat#A-21422 |
| Goat anti-Rabbit IgG (H+L) Cross-Adsorbed Secondary Antibody, Alexa Fluor™ 488 | Thermo Fisher Scientific | Cat#A-11008 |
| Monoclonal Anti-Beta-Actin, Clone AC-74 produced in mouse | MilliporeSigma | Cat#A2228 |
| **Bacterial and Virus Strains** | | |
| Andes virus (Chile-9717869 strain) | Dr. Heinrich Feldmann at NIH/NIAID | NA |
| Sin Nombre virus (SNV-77734) | Dr. Heinrich Feldmann at NIH/NIAID | NA |
| Hantaan virus, Fojnica Strain | BEI Resources | Cat#NR-9370 |
| **Chemicals, Peptides, and Recombinant Proteins** | | |
| Regular Fetal Bovine Serum | Corning | Cat#35010CV |
| Eagle's Minimum Essential Medium (MEM) | Corning | Cat#10009CV |
| Penicillin-Streptomycin (10,000 U/mL) | Gibco | Cat#15140122 |
| L-Glutamine (200 mM) | Gibco | Cat#25030081 |
| MEM Non-Essential Amino Acids Solution(100X) | Gibco | Cat#11140050 |
| Silymarin | MilliporeSigma | Cat#S0292 |
| Urolithin B | Selleckchem | Cat# S1321 |
| Favipiravir (T-705) | Selleckchem | Cat# S7975 |
| 6-Azauridine | MilliporeSigma | Cat#A1882 |
| diABZI STING agonist | Selleckchem | Cat#S8796 |
| cAIMP | Invivogen | Cat#tlrl-nacai |
| GC376 | Selleckchem | Cat#S0475 |
| Cholesterol | MedChemExpress | Cat#HY-N0322 |
| 25-Hydroxycholesterol | MedChemExpress | Cat#HY-113134 |
| RPMI 1640 | Thermo Fisher Scientific | Cat#11875093 |
| B27 supplement with insulin | Thermo Fisher Scientific | Cat#17504044 |
| CHIR-99021 (CT99021) | Selleckchem | Cat#S1263 |
| Recombinant Human IFN-β | Peprotech | Cat#300-02BC |
| Dimethyl sulfoxide | MilliporeSigma | Cat#D2650 |

| | | |
|---|---|---|
| 16% Paraformaldehyde (formaldehyde) aqueous solution | Electron Microscopy Sciences | Cat#15710 |
| Dulbecco's Phosphate-Buffered Salt Solution 1X | Corning | Cat#21030CV |
| DAPI (4',6-Diamidino-2-Phenylindole, Dihydrochloride) | Thermo Fisher Scientific | Cat#D1306 |
| Corning™ Cell Culture Phosphate Buffered Saline (1X) | Thermo Fisher Scientific | Cat#MT21040CV |
| Bovine Serum Albumin | MilliporeSigma | Cat#A9418 |
| Normal Donkey Serum | Jackson ImmunoResearch | Cat#017-000-121 |
| Normal Goat Serum | Cell Signaling | Cat#5425S |
| Triton-X 100 | MilliporeSigma | Cat#T9284 |
| **Commercial Assays** | | |
| CellTiter-Glo Luminescent Cell Viability Assay | Promega | Cat#G7570 |
| **Experimental Models: Cell Lines** | | |
| VERO C1008 [Vero 76, clone E6, Vero E6] | ATCC | Cat#CRL-158 |
| Calu-3 | ATCC | Cat#HTB-55 |
| hPSC derived Astrocytes | Shi laboratory, City of Hope | N/A |
| hPSC derived cardiomyocyte | Deb laboratory, UCLA | N/A |
| Human lung organoid | Stripp laboratory, Cedars Sinai | N/A |
| Mouse lung organoid | Stripp laboratory, Cedars Sinai | N/A |
| **Oligonucleotides** | | |
| Primers for ANDV (Forward: GCTTCTGCTTTCGCATTGC; Reverse:GTGAGGTAGTATGTGTTGAGGTAG) | This Paper | N/A |
| Primers for SNV (Forward: TGACACTAATGCTTGCTTTGC; Reverse: GCAATCAAGAATTTACTTATAATGAGGTAG) | This Paper | N/A |
| Primers for HTNV (Forward: GTTAGTAAGCAGAGAAAGCAGAAAG; Reverse: TGGGTCAGTTAATCCGTTGTG) | This Paper | N/A |
| Primers for Human OAS1 (Forward: TGTGTGTCCAAGGTGGTAAAGGG; Reverse: AAGACAACCAGGTCAGCGTCAG) | This Paper | N/A |
| Primers for Human IFN-λ (Forward: CGCCTTGGAAGAGTCACTCA Reverse: GAAGCCTCAGGTCCCAATTC) | This Paper | N/A |
| Primers for Human IL1-β (Forward: AAGCTGATGGCCCTAAACAG Reverse: AGGTGCATCGTGCACATAAG) | This Paper | N/A |
| Primers for Human IFN-β (Forward: AAGGCCAAGGAGTACAGTC Reverse: ATCTTCAGTTTCGGAGGTAA) | This Paper | N/A |
| **Deposited Data** | | |
| RNA-Seq of Calu-3, hPSC derived cardiomyocytes and hPSC derived Astrocytes | Gene Expression Omnibus | Accession Number: GSE232641 |

| | | | |
|---|---|---|---|
| cells infected with ANDV, HTNV, SNV and/or treated with Silymarin, Urolithin-B, Favipiravir and 6AZA | | | |
| **Software and Algorithms** | | | |
| GraphPad Prism 9 | | GraphPad | N/A |
| Multi-Point Tool (Cell Counter) | | ImageJ | N/A |
| BioRender | | BioRender | N/A |
| RStudio 2023.03.0 | | Posit | N/A |
| **COMPOUND DETAILS** | **MOLECULAR FORMULA/SEQUENCE** | **CAS#** | **STRUCTURE** |
| Silymarin | $C_{25}H_{22}O_{10}$ | 65666-07-1 | 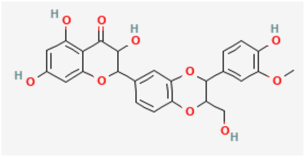 |
| Urolithin B | $C_{13}H_8O_3$ | 1139-83-9 | 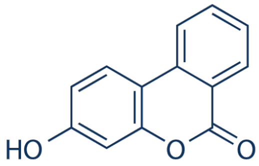 |
| Favipiravir (T-705) | $C_5H_4FN_3O_2$ | 259793-96-9 | 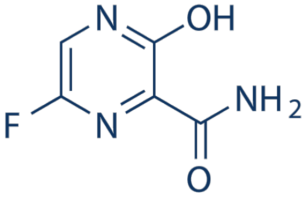 |
| 6-Azauridine | $C_8H_{11}N_3O_6$ | 54-25-1 | 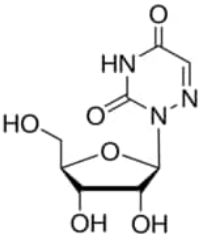 |
| cAIMP | $C_{20}H_{23}N_9O_{13}P_2$ .2Na | 1507367-51-2 | 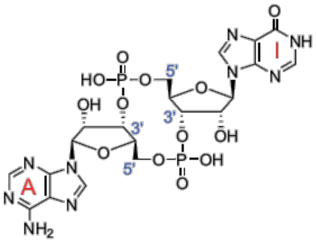 |

| | | | |
|---|---|---|---|
| diABZI STING agonist | $C_{42}H_{51}N_{13}O_7$ | 2138498-18-5 | 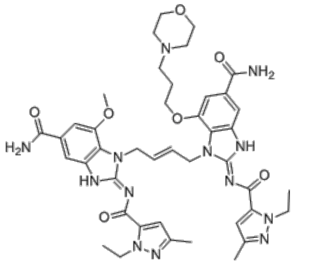 |
| GC376 | $C_{21}H_{30}N_3NaO_8S$ | 1416992-39-6 | 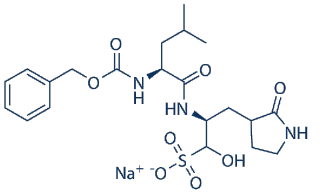 |
| Cholesterol | $C_{27}H_{46}O$ | 57-88-5 | 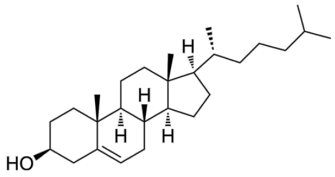 |
| 25-Hydroxycholesterol | $C_{27}H_{46}O_2$ | 2140-46-7 | 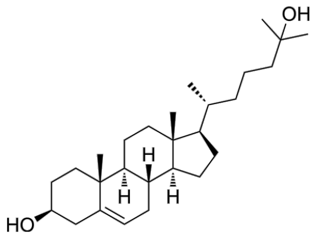 |

**Supplementary Table 5. Reagents and Resources Used in This Study.**

**Supplementary Video 1**. **Comparison of Mock- and ANDV-Infected hPSC-CMs at 48 hpi.** A) Mock-infected hPSC-CMs at 48 hpi. B) ANDV-infected hPSC-CMs at 48 hpi. (Attached separately).
